# Homological landscape of human brain functional sub-circuits

**DOI:** 10.1101/2023.12.22.573062

**Authors:** Duy Duong-Tran, Ralph Kaufmann, Jiong Chen, Xuan Wang, Sumita Garai, Frederick Xu, Jingxuan Bao, Enrico Amico, Alan David Kaplan, Giovanni Petri, Joaquin Goni, Yize Zhao, Li Shen

**Affiliations:** Department of Biostatistics, Epidemiology and Informatics, Perelman School of Medicine, University of Pennsylvania, PA, USA; Department of Mathematics, United States Naval Academy, Annapolis, MD, USA; Department of Mathematics, Purdue University, West Lafayette, IN, USA; Department of Bioengineering, School of Engineering and Applied Science, University of Pennsylvania, PA, USA; Department of Electrical and Computer Engineering, George Mason University, Fairfax, VA, USA; Neuro-X Institute, EPFL, Geneva, Switzerland; Department of Radiology and Medical Informatics, University of Geneva, Switzerland; Computational Engineering Division, Lawrence Livermore National Laboratory, Livermore, CA, USA; CENTAI Institute, 10138 Torino, Italy; NPLab, Network Science Institute, Northeastern University London, London, E1W 1LP, United Kingdom; Networks Unit, IMT Lucca Institute, 55100 Lucca, Italy; Purdue Institute for Integrative Neuroscience, Purdue University, West Lafayette, Indiana, USA; School of Industrial Engineering, Purdue University, West Lafayette, Indiana, USA; Weldon School of Biomedical Engineering, Purdue University, West Lafayette, Indiana, US; School of Public Health, Yale University, New Heaven, CT, USA

## Abstract

Human whole-brain functional connectivity networks have been shown to exhibit both local/quasilocal (e.g., set of functional sub-circuits induced by node or edge attributes) and non-local (e.g., higher-order functional coordination patterns) properties. Nonetheless, the non-local properties of topological strata induced by local/quasilocal functional sub-circuits have yet to be addressed. To that end, we proposed a homological formalism that enables the quantification of higher-order characteristics of human brain functional sub-circuits. Our results indicated that each homological order uniquely unravels diverse, complementary properties of human brain functional sub-circuits. Noticeably, the *H*_1_ homological distance between rest and motor task were observed at both whole-brain and sub-circuit consolidated level which suggested the self-similarity property of human brain functional connectivity unraveled by homological kernel. Furthermore, at the whole-brain level, the rest-task differentiation was found to be most prominent between rest and different tasks at different homological orders: i) Emotion task (*H*_0_), ii) Motor task (*H*_1_), and iii) Working memory task (*H*_2_). At the functional sub-circuit level, the rest-task functional dichotomy of default mode network is found to be mostly prominent at the first and second homological scaffolds. Also at such scale, we found that the limbic network plays a significant role in homological reconfiguration across both task- and subject-domain which sheds light to subsequent investigations on the complex neuro-physiological role of such network. From a wider perspective, our formalism can be applied, beyond brain connectomics, to study non-localized coordination patterns of localized structures stretching across complex network fibers.

## 1 Introduction

Network science sheds light on complex phenomena - from fake news spreading mechanisms in a social network to natural equilibrium in large-scale ecosystems with competing species interactions. Graphs (Networks), despite its convenience and power to unravel many important phenomena from social, financial to biological networks, lack comprehensive ability to describe higher-order dynamics of complex systems [11]. Indeed, many real-world systems, although can be described using diatic relation (edges), have indeed polyadic functionality [35, 36]. Prior studies have strongly suggested the critical role of higher-order interactions in terms of explaining complex intertwined dynamics such as phase transitions of emergent phenomena in networked systems [11]. For instance, higher-order effects emerged from neuronal population are shown to be significant in both statistical, topological, and other domains [36, 56, 60, 77]. Higher order interactions, as formalized by hyperedges (in hypergraphs) or simplicial complexes (in homology), have shown to unravel many complementary functions, compared to node-/edge-based investigations [11].

The human brain is a complex system exhibiting multi-scale property where interactions among its finest elements (e.g., neurons) orchestrate emergent phenomena (e.g., cognition, consciousness [9]). Besides exerting hierarchical cytoarchitecture, human brain functional organizations also display “modular” characteristics - also known as hierarchical modularity [47]. Bullmore and Sporns [17] were among the first investigators noting that whole-brain functional connectivity can be effectively characterized into (functional) modules whose elements (e.g., nodes/vertices in a functional connectome (FC)) are contributed by different distributed areas across the cortex. Specifically, the human brain can be decomposed into specialized, yet highly interactive functional modules [9, 71] (or equivalently, communities in complex networks, see [2, 29, 31] among others). The modular setting of human brain into distinctive functional sub-circuits allows its function to adapt flexibly with diverse cognitive requirements [10,24]. Moreover, functional modularity can also explain human brain complexity [9], cognitive reconfiguration [24], rest-task divergence [5], among other functionalities.

In 2011, the concept of intrinsic functional connectivity Magnetic Resonance Imaging (fcMRI) network (also known as functional sub-circuits, functional network (FN) or resting-state networks (RSNs)) was put forth by Yeo and colleagues [76]. FNs are essentially parallel interdigitated sub-circuits in which each cortical lobe might contain multiple regions belonging to one or more FNs. An *a priori* set of FNs (or equivalently, functional sub-circuits) elucidates different executive functions of human brain in healthy, neurodegenerative disease or developmental conditions [15]. Mathematically, an *a priori* identification of FNs is a partition of the whole-brain functional connectivity which results in a functional atlas (e.g., a guidance to which brain region(s) belong to which functional sub-circuit(s)). Such partition can be used as **a baseline reference** to investigate physiological, functional, individual differences of *i)* the same FN across different cognitive conditions [24] or *ii)* different FNs across the same task (e.g., fMRI). Specifically, the mapping of an *a priori* set of FNs (to different individuals’ functional connectivity) allows the investigation of *i)* the functional differences among individuals under different cognitive demands [24, 45, 49]; *ii)* aging [43, 45, 57]; or *iii)* neurological dysfunctions [26, 46, 75]. Besides Yeo’s functional FN atlas, other highly putative establishments of *a priori* set of FNs also featured Power et al. [51], Glasser et al. [37], Gordon et al. [39], and most recently Schaefer et al. [58]. The most recent review on the identification and applications of *a priori* set of FN mappings can be found in the work of Bryce and colleagues [15].

In the case of human brain complex networks, higher-order interactions among neuron populations, at the whole-brain level, have been shown to unravel complementary insights that otherwise, would not be fully appreciated by conventional node-based (zeroth-order) or edge-based (first-order) investigations [34, 36, 56, 59, 60, 77]. Nonetheless, higher-order characteristics induced from an *a priori* set of FNs have yet been investigated. Understanding complex behaviors arisen at a scale between the mesoscopic (brain regions) and macroscopic (whole-brain) level would set the stage to a deeper, comprehensive picture understanding of the human brain large-scale functional sub-circuitry, which, in turns, provide foundational support to investigate individualized or task-based parcellations [54, 55]. To that end, we formally explored and measured the topological invariant characteristics of an *a priori* set of FNs (e.g., Yeo’s sub-circuitry [76]) through the first three homological dimensions: *H*_0_ (connected components), *H*_1_ (first-order (graph-theoretical) cycles), and *H*_2_ (second-order cycles). These explorations on homological properties of FNs are computed on the 100 unrelated subjects from the Human Connectome Project (HCP) dataset in which fMRI data were recorded, for each subject, in resting state and seven other fMRI tasks. The fMRI data were processed and parcellated into 360 brain regions, according to [38]. To investigate the higher-order mesoscopic properties of the constructed functional connectomes (FCs), we used the seven *a priori* FNs, proposed by Yeo and colleagues [76], the 14 sub-cortical regions are added for completeness. It is worthy to note that our proposed framework can be applied to other combinations of parcellations and functional sub-circuitry partitions.

## 2 Formalism

The progression to glean topological information for a set of data, which by itself is discrete is first turn it into a graph modeling the first-order interactions and then to progress to a topological space by realizing its simplicial clique complex Δ(Γ) which models simultaneous, and thereby higher order, interactions. The topological construction flow is as follows:

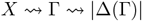

We stress that the first-order information yielding the graph is an additional datum, while the clique complex completes this data to a space. The topological space, which is simplicial in nature, has topological invariants associated to it, such as the homology *H*_*i*_(Δ(Γ)) and Betti numbers *b*_*i*_. The 0th Betti number *b*_0_ counts the number of components and the first Betti number *b*_1_ which counts the number of independent loops (i.e. graph-theoretical cycles). If the graph is connected these satisfy *b*_0_ − *b*_1_ = # of vertices − # of edges. The next higher interaction is *b*_2_ which counts the number of independent spheres, or more precisely homology classes, in the realization. The realization is given by inserting a simplex for each complete graph, see below.

Graphs in this setting are best understood as given by symmetric matrices, the entries of which are given by the first-order interaction as witnessed by Pearson correlation functions. Defining a cut–off parameter *r* for the interactions then determines a graph Γ(*r*) and the homology becomes a function of this *r*. Scanning *r* from 0 to 1 homology is born and annihilated. The sequence of these events is mathematically captured by persistence homology and can be encoded and visualized in terms of bar codes.

When comparing different bar codes, one usually uses the Wasserstein distance, which is a natural norm on the space of such diagram. It is not the only norm though and in special situations other measures are more appropriate.

### 2.1 Graph, induced subgraph, Clique complex

In the context of this study, the graph (network) quantifying whole-brain functional connectivity profile is called the functional connectome (FC). Induced subgraphs are utilized to model functional sub-circuits (e.g., Yeo’s Functional Networks or FNs) of the FC. By construction, an FC is a complete weighted graph. The mathematical and computational setup is as follows:

Mathematically, a *graph/network* Γ with vertex set *V* and edge set of edges *E* where an edge in *E* is a two-element set {*u, v*} of vertices. Enumerating the vertex set by 1, … *n*, a graph is equivalently encoded by its symmetric adjacency matrix *M* (Γ) whose entries are *m*_*uv*_ = 1 if the vertices *u* and *v* are connected by an edge and 0 if not. We make the choice that the diagonal entries are 1. A graph is *complete* if there is an edge between any two distinct nodes. The matrix *M* (Γ) is the matrix all of whose entries 1. The number of edges of a complete graph is 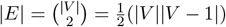 which is the same as the number of non–diagonal independent entries in a symmetric |*V* | *×* |*V* | matrix. The two main topological invariants of a graph are the number of connected components *b*_0_ and the number of loops *b*_1_ = |*E*| − |*V* | + *b*_0_, which are also called the first and second Betti numbers the combination *χ* = *b*_0_ − *b*_1_ = |*V* | − |*E*| is called the Euler-characteristic of the graph.

A *subgraph* is specified by a subset of nodes and a subset of edges connecting these nodes. Each graph is a subgraph of the complete graph on its vertices. This can be thought of as deleting the missing edges from complete graph or equivalently setting the corresponding matrix entries to 0. An *induced subgraph* is simply specified by a subset of vertices. It contains all the edges connecting these vertices. If *V* ^*′*^ is the vertex subset the matrix of the induced subgraph is given by the submatrix *M* (Γ)_*V′V′*_. An induced subgraph is a *clique* if it is itself a complete graph, viz. all the entries of *M* (Γ)_*V′V′*_ are 1.

To use topological or simplicial methods such as homology, one promotes a graph Γ to a simplicial space Δ(Γ). This is not simply the graph itself as glued together from points and intervals, but is more involved. It is the realization of the clique complex. The construction can be understood as an iteration of gluing in simplices. A *n* simplex is the topological space of all vectors (*t*_1_, … *t*_*n*+1_) whose entries are non–negative *t*_*i*_ ≥ 0 and whose sum *t*_1_ + … + *t*_*n*+1_ = 1. The dimension, which is the number of free parameters, is *n*. The gluing procedure starts with the 0 simplices. These are the vertices of Γ viewed as points. In the next step one 1-simplex, which is an interval, is glued in for each edge by identifying the endpoints of the interval with the vertices the edge connects. The higher dimensional simplices are glued in according to complete induced subgraphs. For instance, for any three vertices that are pairwise connected by edges, one glues in a 2–simplex, that is a triangle whose sides are the edges. At the next level one glues in 3–simplices, that is tetrahedra, for each complete graph on 4 verities, which has 6 edges identifying the 4 sides of the tetrahedron with the triangles corresponding to the three edge subsets and so on. The gluing procedure is tantamount to giving the (semi)–simplicial structure which specifies to what the *n* dimension *n* − 1 boundary simplices of an *n* simplex are glued, in such a way that the gluing is consistent with all sub–simplices, regardless of their dimension.

The complete graph on *n* vertices as space realizes to the full *n* simplex. Given an arbitrary graph the realization of the clique complex has such a simplex for each complete induced subgraph and these simplices are glued together by inclusion of subgraphs. This identifies the simplex of a subgraph of a complete graph as a side of the simplex of the graph and hence the space is glued together from maximal simplices corresponding to maximal complete subgraphs along faces corresponding to common subgraphs. One can iteratively construct this space by gluing in higher and higher simplices. This space is higher dimensional and has more topologicial invariants, the higher Betti numbers *b*_*i*_ which are the dimensions or ranks of the respective homology groups *H*_*i*_. The number of connected components is the same for the graph and the associated space. The first Betti number *b*_1_ may differ depending on whether one is looking at the graph or the space. The first graph Betti number for the complete graph is 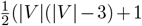 while the first Betti number of the corresponding space, the simplex, is 0.

### 2.2 Filtration by weights and persistent homology

#### Preface

A *non-negatively weighted graph* is a graph together with a weight function *w* : *E* → [0, 1] on its edges. Again, after enumerating the vertices this defines a symmetric matrix *W* = *W* (Γ, *w*) with entries *w*_*uv*_ = *w*({*u, v*}), i.e. the weight of the edge connecting *u* and *v*. If there is no such edge the entry is 0, and the diagonal entries are fixed to be 1. Choosing a cut–off *r* defines the symmetric matrix *W* (*r*) whose entry *w*(*r*)_*uv*_ = 1 if *w*_*uv*_ ≥ *r* and 0 if *w*_*uv*_ *< r*. It has 1’s on the diagonal and defines the graph Γ(*r*). Note that Γ(0) is the original graph and Γ(1) is the graph on the vertex set with no edges. Let 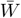 be the order set containing unique weight values, in decreasing order, in matrix *W*, varying the threshold parameter *r* from 0 to 1, defines a sequence of subgraphs as follows:

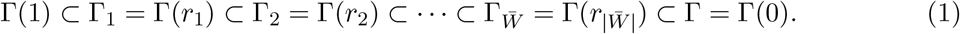

with 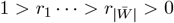 and the Γ_*j*_ are the finitely many different graphs that appear. At each stage *j* some edges are added from the lower stage *j* − 1. The graph Γ(1) is the subgraph with full vertex set, whose edges are given by the non–diagonal entries 1. In practice, if the weights are Pearson correlations functions, the only entries of 1 will be along the diagonal and the graph Γ(1) is simply the discrete set of data.

Note that since set *W* describes diatic **functional couplings** (e.g., similarity) between two nodes of a network (or brain regions of interest (ROIs) in this formalism), it implies that the “distance” (e.g., dissimilarity) between two nodes is defined as follows:

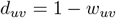

In other words, with this setup, we ensure that

- Γ_1_ is essential the 1-skeleton scaffold where all nodes are perfectly coupled (*d*_*uv*_ = 0), which results in an empty graph.
- 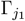 is always an induced subgraph of 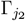 for all 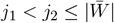;
- The sequence 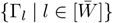 starts with an empty graph (homeomorphic to ℤ_*n*_) and ends with a complete graph (or a clique of *n* nodes) (homeomorphic to simplicial complex of size *n*, i.e. *K*_*n*_).

Given a filtered system, that is a sequence of inclusions of spaces as (1), one can utilize the tool of persistence homology to track the changes of the fundamental topological invariants of homology and Betti-numbers. This supplies a characteristic for the whole sequence. We wish to stress that it is the sequence that is of importance here. The two endpoints have rather trivial topological properties. If the start is just the data, then this is a discrete set, and at the other end the space is just a full simplex corresponding to the complete graph, which is contractible. The transition from one to the other and the appearance —and disappearance— of higher homology is what is kept track of by persistent homology.

#### Bar codes and distances between them

The fingerprint is the variation which is quantified by the bar codes. The variation parameter is the parameter *r* introduced above. A bar code is a type of signature for the variation. For each persistent homology class it records the value of the parameter *r*_*ini*_ when a representative appears (birth) and the value *r*_*fin*_ when it disappears (death). This is an interval (or bar) [*b*(*c*) = *r*_*int*_, *d(c*) = *r*_*fin*_]. At any given *r* the homology is given by those classes *c* for which *r* ∈ [*b*(*c*), *d*(*c*)]. In the variation all higher homology classes are born and eventually die. The 0-th homology starts with as many classes as data points and then eventually decreases (classes die) until there is only one class left, which says that space is connected. The bar code is equivalently encoded by the persistence diagram which the set with multiplicity (multiset) of all the endpoints of the bars {(*b*(*c*), *d*(*c*))}. This is actually a multi–set, since some of the classes may appear and die at the same parameter values and these multiplicities are recorded, e.g. (.2, .8) with multiplicity 2 means that there are two bars of this type. Its *p*–th part *Dgm*_*p*_, is given by bar corresponding to classes of homological dimension *p*.

#### Topological distance formulation

The Wasserstein distance is the natural norm on the diagram space, e.g. the birth-death diagram of topological features. The Wasserstein distance is the right measure for processes taking one diagram to another in a varying family —now of persistence diagrams. This is well suited for analyzing a basic underlying setup with variations. This is commonly viewed and addressed as the stability theorem. In our case, we use Wasserstein distance to compute the distance between two diagrams for the first and second-order homology (e.g., *p* = {1, 2}) in various scenarios (e.g., comparing topological behaviors between the same functional networks at resting conditions).

Specifically, for a fixed homological order *p* (in this paper, *p* = {1, 2}), the *q*−Wasserstein distance *D*_*W,q*_ (∀*q >* 1) for two persistent diagrams *Dgm*_*p*_(*X*) and *Dgm*_*p*_(*Y*) for two data sets *X, Y* can be defined as follows [18]. For a single interval *I* = [*x, y*] set 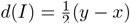 which is the distance to the diagonal of the point (*x, y*) in ℝ^2^. For two intervals *I* = [*x*_1_, *y*_1_], *J* = [*x*_2_, *y*_2_] define their distance as *d*(*I, J*) = max(|*x*_2_ − *x*_1_|, |*y*_2_ − *y*_1_|). This is the max norm distance for the two points (*x*_1_, *y*_1_), (*x*_2_, *y*_2_) in ℝ^2^. A partial pairings between two sets *S* and *T* is a choice of subsets *S*_0_ ⊂ *S, T*_0_ ⊂ *T* and a 1-1 correspondence between the two subsets *π* : *S*_0_ ⇔ *T*_0_. This extends to sets with multiplicity by choosing multiplicities of elements and matching them with multiplicity. Given diagrams *Dgm*_*p*_(*X*), *Dgm*_*p*_(*Y*) let Π be the set of all partial pairings then. The Wasserstein distance minimizes the sum of three contributions: the distances between intervals that are paired and two contributions of the distance to the diagonal for intervals that are not paired. It minimizes over two possible scenarios, points moving and points moving in and out of the diagonal. The first means that the classes shift in their rates and the second means that the classes vanish from the diagram and new classes are introduced. Given *π* let *Dgm*_*p*_(*X*)_1_ = *Dgm*_*p*_(*X*) \ *Dgm*_*p*_(*X*)_0_ and *Dgm*_*p*_(*Y*)_=_*Dgm*_*p*_(*Y*) \ *Dgm*_*p*_(*Y*)_0_)^*q*^ be the complements.

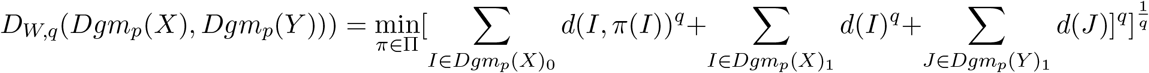

In the zeroth order homology the Wasserstein distance becomes an unnatural choice. This is due to the fact that the data points are the 0–classes and they are all born at *r* = 0. Thus a contribution as disappearing or appearing from the diagonal which signifies being born at different times is not a possible scenario.

It is better to consider *Dgm*_0_(*X*) just as the multiset of endpoints of the bars [0, *d*(*x*)] where *x* ∈ *X* and use the classical Hausdorff distance to measure the (dis-)similarity between two point clouds living in ℝ. This specialized to:

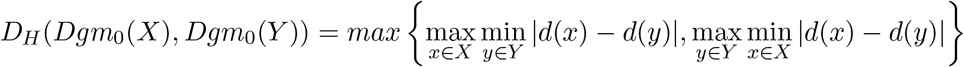

### 2.3 Functional connectomes and mesoscopic structures

Mesoscopic structures are typically referred to as structures whose elements are proper subsets of system’s elements. In brain connectomics domain, there are two types of mesoscopic structures: localized/quasilocalized and non-localized (topological strata). In this section, we provide an overview and definition of each type in the context of brain connectivity.

#### 2.3.1 Localized mesoscopic structures

Localized mesoscopic structures are sub-systems that are learned from local network properties such as nodes or edges, or correlations among neighboring nodes. In brain connectomics, these sub-structures are induced from a wide array of techniques, including but not limited to clustering [51, 76], low dimensional approximation of high-dimensional dynamics [62–66]. The most commonly known localized mesoscopic structures in brain networks are often referred to as functional sub-circuits or functional networks [71].

##### Definition 1

*(Definition adapted from [22]) An a priori set of Functional networks (FNs) are sub-circuits (or equivalently, sub-networks) that are highly reproducible across individuals at resting condition (absence of task-induced cognitive demand). Hence, FNs are also known as Resting-State networks (RSNs)*.

Special collections of induced subgraphs are used to group brain regions of interest (ROIs) into localized/quasilocalized mesoscopic structures of brain functions denoted as functional sub-circuits or equivalently, functional networks (FNs). A collection of *k* subgraphs (of graph Γ) is denoted as {*γ*_*i*_ ⊂ Γ | *i* ∈ [*k*]}. A collection of induced subgraphs is *a vertex covering* if the graphs each vertex of Γ is a vertex of one of the *γ*_*i*_. Such a vertex covering is disjoint if the *γ*_*i*_ have disjoint vertices. After enumerating all nodes by 1, …, *n* = |*V* | the collection of induced subgraphs is fixed by the membership assignment. This is specified by a partition vector denoted as *σ* ∈ [*k*]^*n*^ where *σ* = [*σ*_*u*_] = *i* ∈ [*k*] indicating that *u* belongs to *γ*_*i*_ | *i* = {1, 2, …, *k*}. Note that in network science, FNs are equivalent to the term “communities” [2, 3, 29, 31]. The problem of identifying the set of communities {*γ*_*i*_ ⊂ Γ | *i* ∈ [*k*]} for a given complex network is called the community detection problem [2, 3, 29–31].

#### 2.3.2 Non-localized mesoscopic structures

While studies of network properties and dynamics using locally featured properties (nodes, edges attributes) provided a well-grounded approach, these methods were proven to be cumbersome in describing and quantifying heterogeneity existing across network dynamical fabrics. These structures usually encompass many-body interactions or encapsulate topological sub-structures that can not be mathematically described using local attributes. To that end, homology [25] offers a unique capability to capture the so-called non-localized mesoscopic structures that otherwise, cannot be reduced to local or quasilocal network properties. In the context of weighted complex networks, persistent homology is used to identify how long (the persistence of) a hole (at any given dimension) lasts from its birth (the weight scale 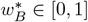 that the hole is observed) to its death (the weight scale 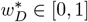 that the hole is filled).

In the context of functional brain connectivity, non-localized mesoscopic structures in a FC represent the encapsulated area where there is *less* functional connectivity collectively formed among brain regions encapsulating these structures [50]. Such structure characterizes the notion of hole; the boundary that wraps around these structures are the non-localized mesoscopic fabrics characterized by the so-called cycles. These cycles exist in different homological dimensions for a given networked system which can be described in the language of a manifold. The hollow structures (holes) could be seen as overarching wraps-around special hollow structures in a manifold with different characteristics and properties, compared to functional networks [13, 23, 27, 76] or communities [2, 3, 28, 29] in complex networks.

### 2.4 Consolidated/Super graph

The system under consideration is naturally regarded as a two–level system given by the ROIs and their connections. The first level is made up of the individual ROIs and the second level is given by the connections between the ROIs. In graph theoretical language, the full graph Γ(*r*) containing all the nodes naturally has a subgraph *γ*_*i*_(*r*) ⊂ Γ(*r*). These subgraphs form a supergraph, which has the subgraph as new vertices and has the edges between two vertices if there are edges between the subgraphs. There are two versions, the first is the multi–edged graph that is described theoretically by contracting all the edges of the subgraphs *γ*_*i*_ that is if *γ* = ∪_*i*_*γ*_*i*_ is the union of subgraphs, then 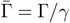. Reducing possible multiple edges to just one edge on has the reduced graph 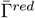 which is again an ordinary graph described by a matrix. For a weighted graph, assuming the subgraphs are not connected, the graphs *γ*_*i*_ correspond to block matrices along the diagonal and the edges of the quotient graph are the off block entries. To obtain a matrix one can consolidate the weights into one weight by choosing a function 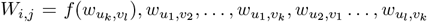 where *u*_1_, … *u*_*l*_ are the vertices of *γ*_*i*_ and *v*_1_, …, *v*_*l*_ ar the vertices of *γ*_*i*_. One such choice is 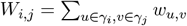 and then normalize to

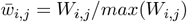

In the case under consideration the graph 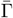 has eight (super-)vertices corresponding to each FNs. The basic topological invariant of the loop number is of great interest as it is a measure of the inter-connectivity of these “super-regions”. The persistent homology for the normalized super graph, that is the consolidated graph, will then complement this information to show clusters of correlations between FNs.

## 3 Results

### 3.1 Data

#### Human Connectome Project (HCP) Dataset

We used the master data release extracted from the HCP Young Adult (HCP-YA) subject release [74]. Specifically, the fMRI dataset is obtained from HCP repository (http://www.humanconnectome.org/), with Released Q3. In general, all MRI neuroimaging modalities were acquired in two different days, with two different scanning patterns (e.g., phase acquisitions: left to right or LR and right-to-left or RL). The detailed description is in the next section and **Figure** 2.

**Figure 1.**
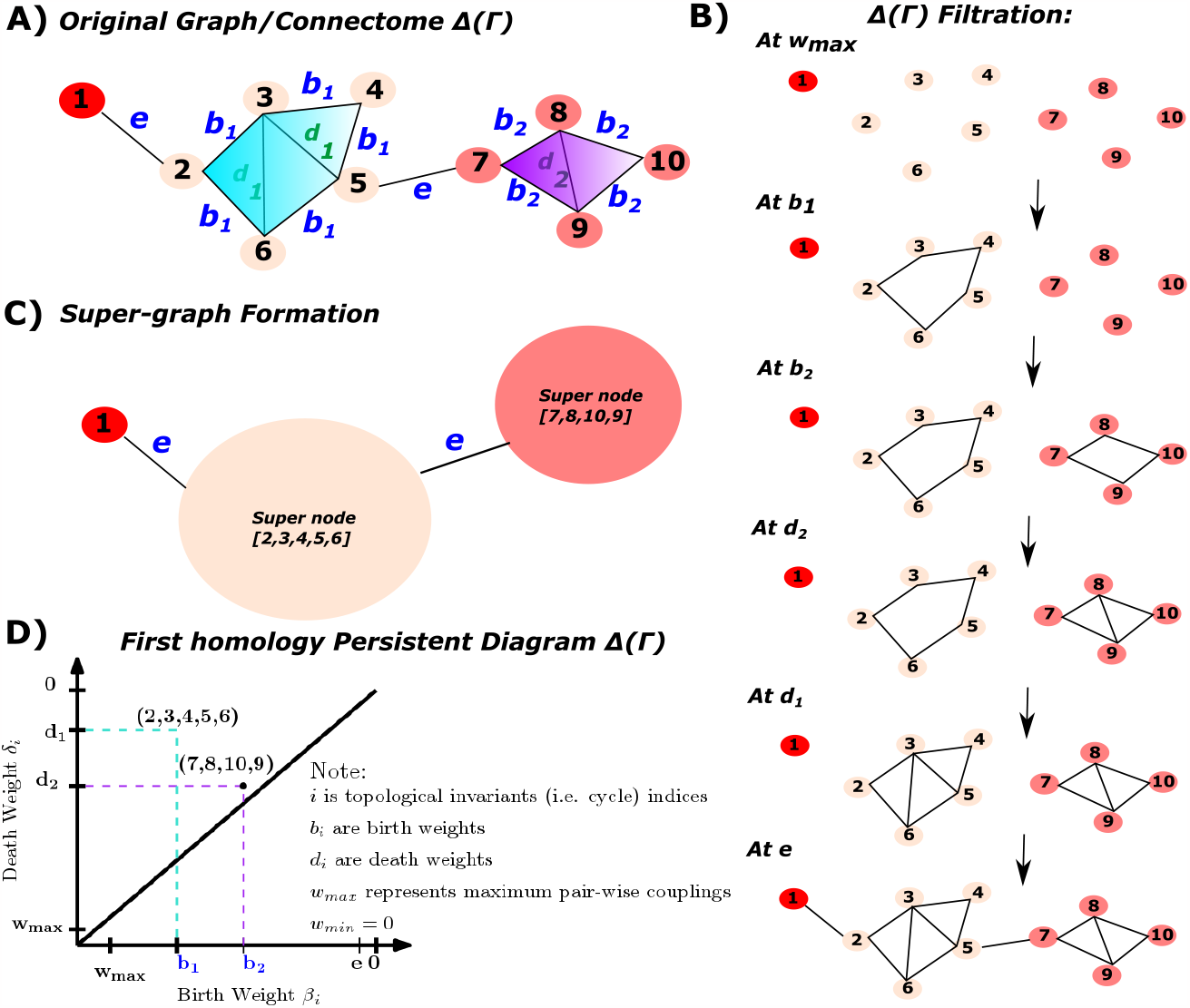
Topological landscape of human brain functional networks: **Panel A** is the schematic representation of a graph (e.g., functional connectome) modeling first-order interactions (e.g., functional couplings) with weight values *w*_*uv*_ = {*d*_1_, *d*_2_, *e, b*_2_, *b*_1_}. **Panel B** is a sequence of induced subgraph scaffolds (also referred to as filtration) by scanning across *w*_*uv*_ (Note that the filtration is built on *d*_*uv*_ = 1 − *w*_*uv*_); hence, the starting point Γ(*w*_*max*_ = 1) = Γ_1_ is an empty graph. **Panel C** represents the super-graph construction by merging all ROIs belong to the same FN to one super-node through equivalence relation 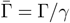 which is defined as follows: *γ*_1_ = {1} (e.g., FN1); *γ*_2_ = {2, 3, 4, 5, 6} (e.g., FN2) and *γ*_3_ = {7, 8, 9, 10} (e.g., FN3). Notice that the super-graph itself is a graph; hence, homological computations that were applied in the original graph can also be applied to the super-graph itself. In this example, the super-/consolidated graph has 3 super-nodes. Additionally, the weight matrix is re-scaled according to 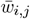. **Panel D** is the corresponding persistent diagram for the first homology which accounts for two first-order cycles in a network: **(2**,**3**,**4**,**5**,**6)** and **(7**,**8**,**10**,**9)**; here, we see that cycle **(2**,**3**,**4**,**5**,**6)** lasts longer (more persistent) compared to cycle **(7**,**8**,**10**,**9)**. Finally, when scanning across five distinct *r* parameters, we obtain the zeroth and first Betti numbers: *b*_0_ = {10, 6, 3, 3, 3, 1} and *b*_1_ = {0, 1, 2, 1, 0, 0}, respectively.

**Figure 2.**
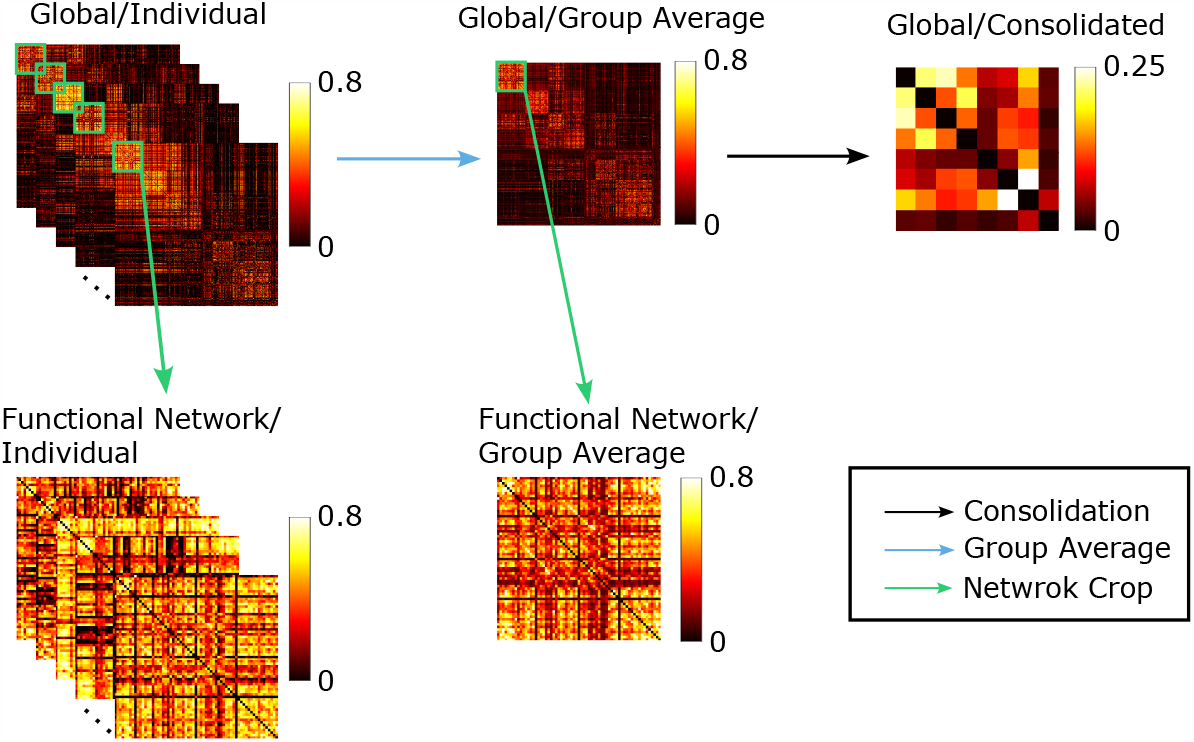
fMRI whole-brain connectome multi-level analysis workflow. For each task, we started with individual-level functional connectome. On the global (macroscopic) level, we have individual analysis as well as group-averaged analysis, and the functional network (mesoscopic) level extracts functional networks from either the individual or group-averaged macroscopic graph. The consolidated graph is constructed by aggregating the nodes from the group-averaged macroscopic level connectome.

#### HCP Functional Data

The fMRI data from the 100 unrelated subjects in the HCP Q3 release were employed in this study [73, 74]. Following the HCP protocol, all subjects had provided written consent to the HCP consortium. The two resting-state functional MRI acquisitions with HCP filenames: *rf MRI_REST*_1_ and *rf MRI_REST*_2_ were collected in two separate sessions (on two different days), with two distinct scanning acquisitions (LR and RL) for each day, see [38], [74], and [73] for further details. Besides resting state, the dataset also includes fMRI data from seven (07) fMRI tasks: gambling (*tf MRI_GAMBLING*), relational or reasoning (*tf MRI_RELATIONAL*), social (*tf MRI_SOCIAL*), working memory (*tf MRI_WM*), motor (*tf MRI_MOTOR*), language (*tf MRI_LANGUAGE*), and emotion (*tf MRI_EMOTION*). Per [38], [8], three following fMRI tasks were obtained on the first day: working memory, motor, and gambling; the rest were obtained on the second day. The local Institutional Review Board at Washington University in St. Louis (scan site) approves all the scanning protocols used during the HCP dataset acquisition process used in this paper. Please refer to [8, 38, 68] for a further detailed description of the HCP-YA dataset. Note that all tasks and resting functional MRIs are treated with equal importance. In this work, we denote seven fMRI tasks as gambling (GAM), relational (REL), social (SOC), working memory (WM), language processing (LANG), emotion (EMOT), and motor (MOT).

Table 1 depicts basic information about fMRI conditions’ run time and the number of time points for each task. Subsequently, along with table 1, a brief description of each fMRI condition is provided below. An extended description is provided in HCP manual^1^.

**Table 1.** fMRI task scanning length and number of frames description. All fMRI task run times were reported in order of minutes and seconds. Except for the resting state (for which, each subject was scanned twice per day for a total of 2 *×* 2 = 4 sessions), all other tasks have two scans (RL and LR). TR is the time between two consecutive readings.

| fMRI Conditions | Run time (min:sec) | # of time points |
| --- | --- | --- |
| REST1 (& REST2) | 14:33 | 1200 |
| EMOTION (EMOT) | 2:16 | 176 |
| GAMBLING (GAM) | 3:12 | 253 |
| MOTOR (MOT) | 3:34 | 284 |
| LANGUAGE (LANG) | 3:57 | 316 |
| RELATIONAL (REL) | 2:56 | 232 |
| SOCIAL (SOC) | 3:27 | 274 |
| WORKING MEMORY (WM) | 5:01 | 405 |

1. **REST**: Eye open with relaxed fixation on a bright cross-hair with dark background. 1200 time points were obtained with 720 ms TR.
2. **EMOTION**: Subject was instructed to match two faces (or shapes) are shown at the bottom to the top of the screen. Faces are shown with angry/fearful expression. Each scan involves 3 face blocks and 3 shape blocks with 8 seconds of fixation.
3. **GAMBLING**: card playing game where subject needed to guess a number of a card in order to win or lose money. At each trial, subject was instructed to guess whether a card has value larger or smaller than 5, given the numerical range of the cards was between 1 and 9. Subjects had 1.5 seconds to respond and 1 second of feedback.
4. **LANGUAGE**: At each scan, four blocks of story tasks and four blocks of math tasks were presented to the subject. Stories contained brief auditory information followed by choice of questions about the story topics. Math tasks contained arithmetic questions with a similar level of difficulty compared to the story task.
5. **MOTOR**: subjects were shown various cues and instructed to either tap (left and right) fingers, squeeze (left or right) toes, or move tongue in response to different areas of human brain motor cortex. The task contains a total of 10 movements (12 seconds per movement), preceded by a 3-second cue.
6. **RELATIONAL**: subject were presented with 6 shapes along with 6 different textures. Given two pairs of objects (one on the top and the other one at the bottom of the screen), the subject had to decide whether the shape (or texture) differed across the pair on the top screen. In addition, they had to decide whether the same difference got carried over the bottom pair.
7. **SOCIAL**: subjects are shown a 20-second video clip containing randomly moving objects of various geometrical shapes (squares, circles, triangles, etc.). After that, the subject was instructed to respond whether these objects had any mental interactions (shapes took into account feelings and thoughts), Undecided, or No Interactions.
8. **WORKING MEMORY**: subject was presented with trials of tools, faces, and body parts. Four different stimulus types were presented in each run. In addition, at each run, two types of memory tasks were presented: two-back and zero-back memory tasks.

#### Brain atlas

The brain atlas used in this work is based on the cortical parcellation of 360 brain regions proposed by Glasser and colleagues [37]. Similarly to the description in [6, 7, 24], 14 sub-cortical regions were added for completeness, as provided by the HCP release (filename *Atlas ROI*2.*nii*.*gz*). We accomplish this by converting this file from NIFTI to CIFTI format, using the HCP workbench software^2^ through the command -cifti-create-label. We then obtained a brain atlas of 374 brain regions (360 cortical + 14 sub-cortical nodes) registered to a common space which allowed us to parcellate fMRI voxel-level BOLD time series into brain region of interest level time series (command: -cifti-parcellate). Time series were z-scored by using command -cifti-math.

#### Estimation of functional connectomes

Parcellated time-series were then used to construct the whole-brain functional connectivity by computing the Pearson’s correlation coefficients for each pair of brain regions. This operation can be completed using *Matlab* command -corr which results in a symmetric matrix. All entries in the whole-brain FCs were applied the absolute values so that the threshold parameter *r* = [0, 1].

#### The mapping of functional networks onto FCs

After each subject is registered to the appropriate common space and properly parcellated according to Glasser’s parcellation, we explore the topological features of human brain functional connectivity (FC) by further subdividing whole-brain FC into Resting State Networks (equivalently referred to as functional networks/communities), see [76]. This particular partition includes seven functional networks (FNs): Visual (VIS), SomatoMotor (SM), Dorsal Attention (DA), Ventral Attention (VA), Limbic (LIM), Frontoparietal (FP), Default Mode Network (DMN); Sub-cortical (SUBC) region is, as mentioned above, added into this atlas for completeness. Consequently, the parcellation comprised of eight (8) FNs for each subject/task.

### 3.2 Group analysis: Macroscopic whole-brain Level

#### Topological differences between rest and fMRI tasks

We first explore the topological distances at the group-average whole-brain connectivity level between resting state and fMRI task activation states (see **Figure** 3, see **Figure** S1 for the persistent diagram at the macroscopic level). Each homological group consists of three figures, the first one is the bottom left heatmap, representing the pair-wise Wasserstein distance. The bottom right bar plots show the average distance between one task to all other tasks, thus the task with the highest average distance will indicate its high differentiation with other tasks. Finally, the top right plot shows the variance of each task looking at their distance from the other tasks. Specifically, the zeroth homology suggests that the relational task is the most different to the emotion task. Indeed, other studies, such as [24] through network morphospace mechanism, have also suggested that relational and emotion tasks activate minimally-to-none overlapping functional circuits of the human brain. In terms of *H*_0_ (i.e. connected components), relational task is also the most distinctive task, compared with others (highest average); relational task is followed by resting state on average difference with other tasks. Moreover (see **Figure** 3B), the first homology exhibits the highest degree of differentiation between resting state and task-positive state, as measured by average first homological Wasserstein distance between rest and task bar codes. The first homology also suggests that the motor task is the most topologically different task, compared to the resting state. This finding was consistent with current literature (e.g., Amico and colleague [5]) which stated that motor task exhibited the most distant “within-functional network” edges, relative to other fMRI tasks in the HCP dataset. This result also suggests that at a global scale, the motor cortex whose brain regions are largely employed by motor task, modulates increasing functional activities through forming global transduction pathways with “loop-like” feedbacks (e.g., first-order cycles).

**Figure 3.**
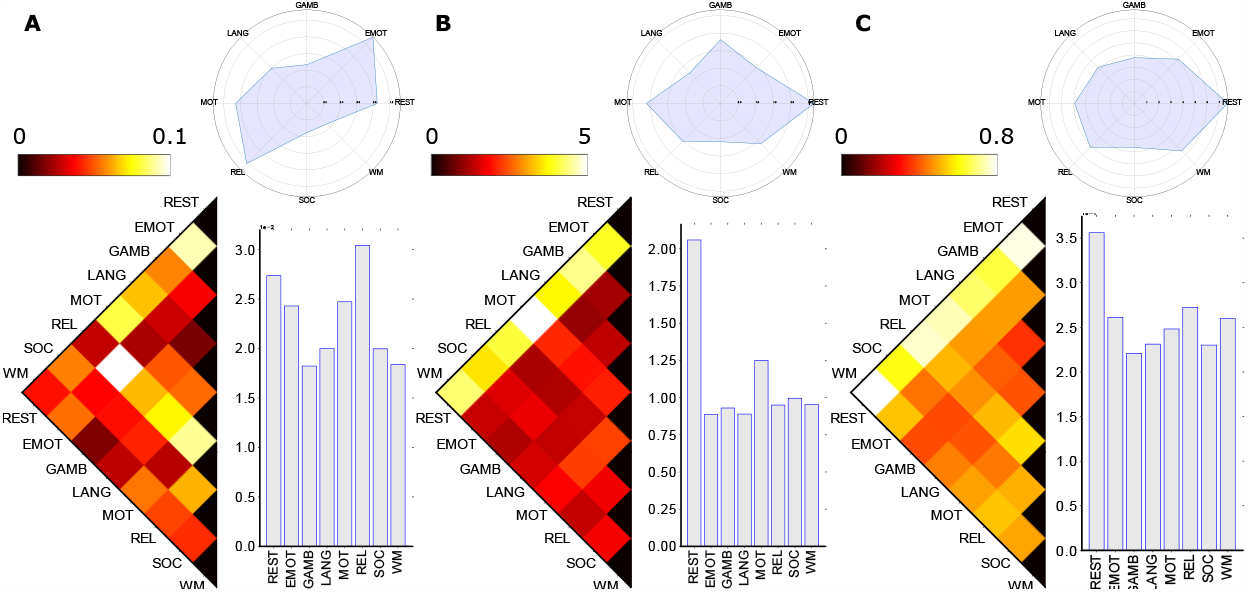
Group-Average Macroscopic Homological distances between fMRI tasks and rest. Specifically, three panels (e.g., left, middle, and right) represent the zeroth (**Panel A**), first (**Panel B**), and second (**Panel C**) homological distance respectively between fMRI tasks and resting condition. Group-average FCs are computed by taking the average of all subjects in the 100 unrelated subjects dataset sampled from the HCP project. The zeroth homological distance is computed using the Hausdorff formula (measured between persistent diagrams of two FNs extracted from group average FC) while the first and second homology distances are computed using the Wasserstein formula. In each panel, the left triangular heatmap represents the distance; the bar plots represent the average distance; and the circular plots represent the variance among fMRI tasks.

### 3.3 Group analysis: Consolidated graph 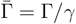

With the construction of consolidated graphs, we generated a smaller-scale representation of the brain connectome to 8 super nodes, which includes 7 Yeo functional networks and one node for subcortical regions. Here, the super-graph is constructed using the equivalence relation at the node level. As such: 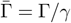 such that *γ* = {*u* ∼ *v* | *σ*_*u*_ = *σ*_*v*_, ∀*u, v* ∈ *V* }. In other words, all brain ROIs belong to the same functional network are contractible.

Since the graph is much smaller, no birth was detected for a 2D simplicial complex in the filtration process, thus only zeroth and first homology were included in the analysis (see **Figure** 4, see **Figure** S2 for the persistent diagram at the consolidated level). In the consolidated setting, we found that the social-resting task pair has the highest distance with the zeroth homology, indicating that in the Yeo functional network level, the connectivity representation captured more differences in social task and resting states (see **Figure** 4A). By the nature of zeroth homology, where we are looking at connected components, the different most-distinct task pair between the global level and consolidated level indicates the choice of representation could impact the topological configuration in brain connectivity. However, the Wasserstein distance between different tasks in the first homology revealed topological invariant among both the global scale as well as node-aggregation scale as the resting state and motor task pair also have the highest distance measure (see **Figure** 4B). This consistency validated the robustness of the first persistent homology class in disentangling the brain’s functional circuits. In addition to the consistency in the most distinct task pair, the resting state task also consistently appears as the most differentiated task compared to other tasks based on the average distance for each task [21, 42]. This indicates that there is a significant reorganization in brain connectivity when people engage in activities from a resting state. Especially for motor tasks, it engages more different brain regions than other tasks, and thus it is also the second distinct task as it is the task that requires responses involving movement.

**Figure 4.**
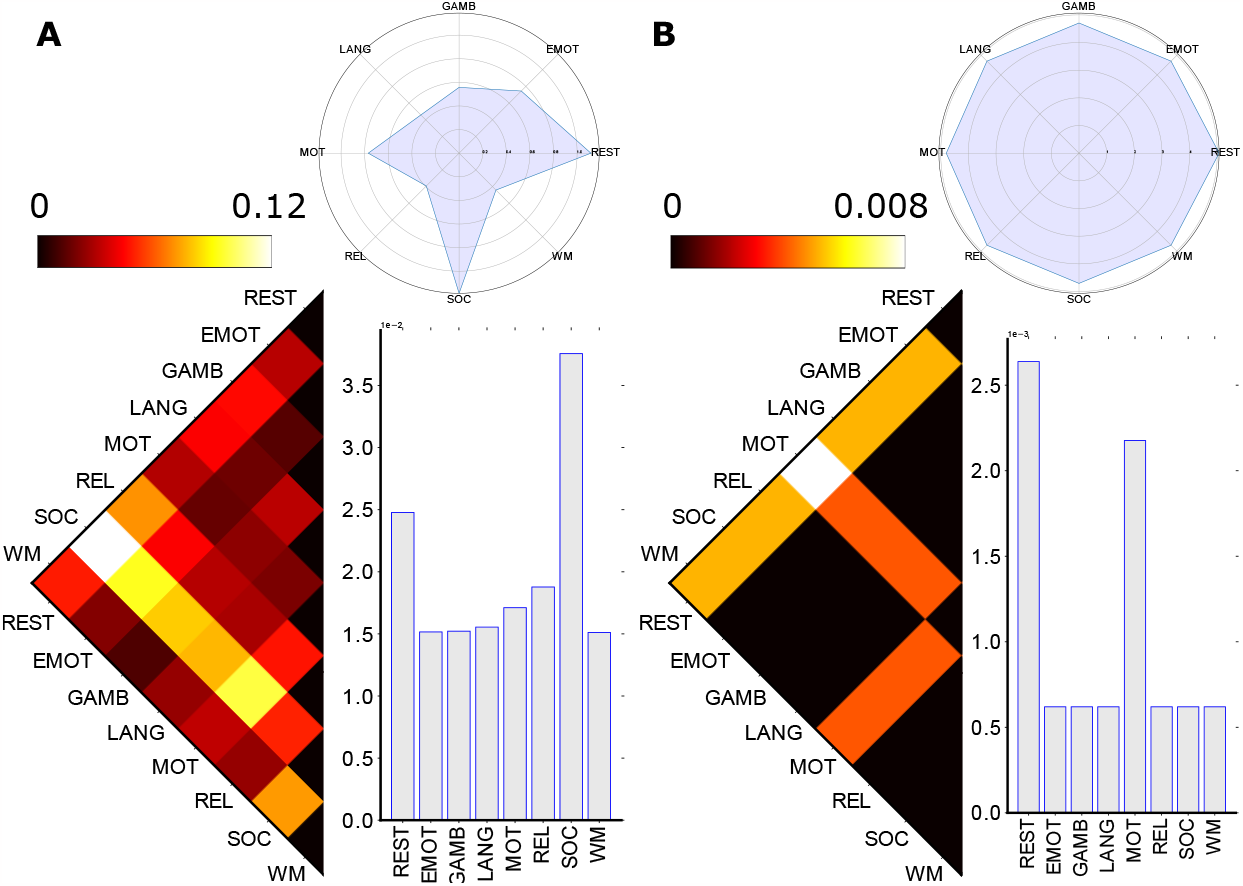
Consolidated homological distances between fMRI tasks and rest. The left and right panel represents the distance between tasks in zeroth (**Panel A**) and first (**Panel B**) homology, calculated by Hausdorff distance and Wasserstein distance respectively. Each panel also contains three components, including the task-wise distance, the average distance, and the variance plot. Due to the small size of the consolidated graph, there was no second homology detected in the corresponding topological space.

### 3.4 Group analysis: Functional network (mesoscopic) Level

In previous sections, we calculated the Wasserstein distance between different tasks, where all of the nodes in the brain connectome were included. In order to assess for a given task, how the brain connectivity shifts from one functional network to another, we also conducted mesoscopic level analysis by extracting the 8 functional networks from the group-averaged global graph. Since previous discoveries showed that the resting state task involves brain regions that are most distinct from other tasks, and the Yeo functional network was also optimized on the resting state fMRI, we focused our analysis on the distance between functional networks in the resting state task and the mesoscopic level topological configuration (see **Figure** S3 - S9 for the analysis of remaining 7 tasks).

#### 3.4.1 Resting state analysis

Fixing the task and extracting functional networks enabled the characterization of within-brain connectivity and the identification of unique topological patterns in functional networks. Particularly, the default mode is present in the pair with the largest Wasserstein distance in H0, H1, and H2 homology, and it also has the largest average Wasserstein distance in H1 and H2 analysis (see **Figure** 5), suggesting a significant level of functional specialization within the default mode during resting state. Extensive studies and literature have validated that the default mode is more active and involved in introspective processes and is typically deactivated in the engagement of goal-oriented tasks, which is referred to as the “resting state dichotomy” of default mode network [16, 33, 52]. This finding further reassured the robustness of the capability of the topological system to detect unique features in certain activities.

**Figure 5.**
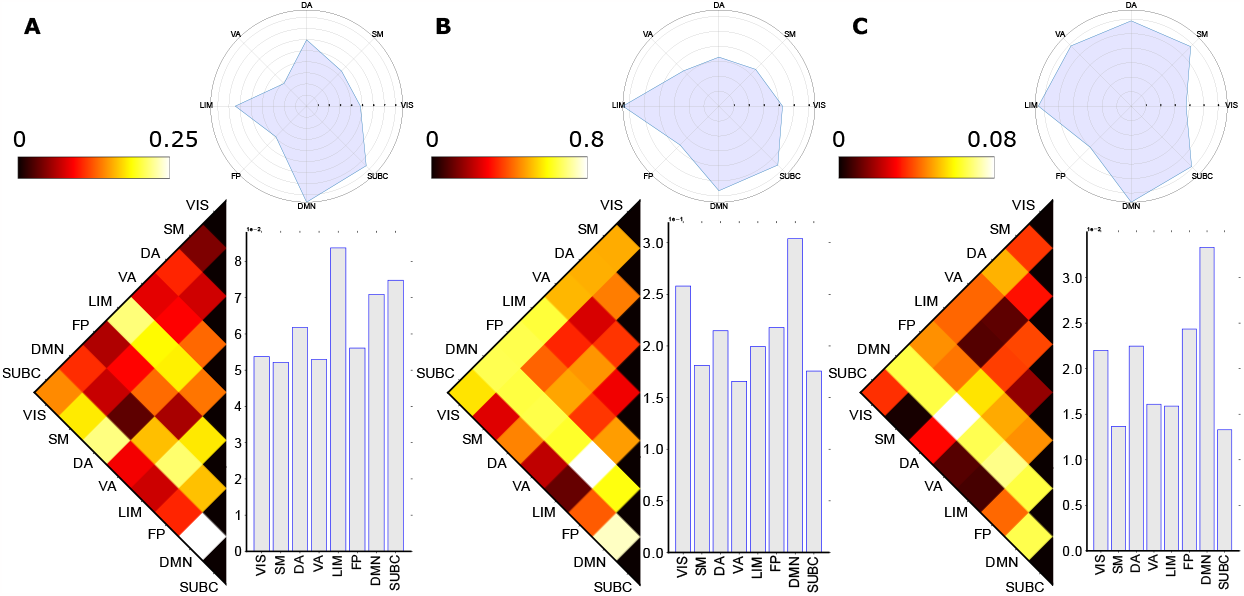
Group-Average homological distances between brain circuits (FNs) at rest (e.g. Resting State Networks). Three panels are positioned similarly to previous figures where they represent the distance of zeroth homology (**Panel A**), first homology (**Panel B**), and second homology (**Panel C**) between pairs of FNs. Group-average FNs are extracted based on Yeo’s parcellation. The zeroth homological distance is computed using the Hausdorff formula while the first and second homology distance are computed using the Wasserstein formula. Each panel contains the triangular distance heatmap, the average distance bar plot, and the variance circular plots among functional networks.

In addition, we also discovered that the limbic system has the highest average Wasserstein distance in the zeroth homology, indicating that it is the most distinct functional network when we compare the pattern of connected components between functional networks [5, 78] (see **Figure** 5A). The limbic system is known for its role in memory- and emotion-related activities [19, 41, 53], and the distinct connectivity pattern discovered reveals that there might still be some memory or emotional processing even during the resting state. Furthermore, the results can also serve as an indication of the individual heterogeneity in their resting state behavior which may involve slight mind activities. The high level of differentiation in H0 task pair with the limbic system is also reconfirmed in the mesoscopic level analysis in emotion task and working memory task (see **Figure** S3A and S9A)

### 3.5 Individual subject analysis

While the group-average level connectomes (global level, consolidated level, and mesoscopic level) provide topological insights in a collective pattern, transitioning to the individual level could further offer a more personalized perspective with after-persistent-homology group insights. Moving beyond the aggregation of group data, individual-level analysis would also allow the consideration of inter-individual variability and consistency across different scales to bring even more robustness to the experimental design. Similar to the previous setting, we investigated the individual global level with consensus voting as well as the individual mesoscopic level with Kullback–Leibler divergence (KL divergence) respectively [44].

#### 3.5.1 Macroscopic whole-brain level

With 100 unrelated subjects from the HCP database, the individual macroscopic level analysis contains 100 independent persistent homology with pair-wise task distance. At the individual macroscopic analysis, we still used the Hausdorff distance for the zeroth homology and the Wasserstein distance for the first and second homology. We evaluated the most distinct pair of tasks in each individual and **Figure** 6 shows the number of times each pair of tasks appeared as the most differentiated task pair.

**Figure 6.**
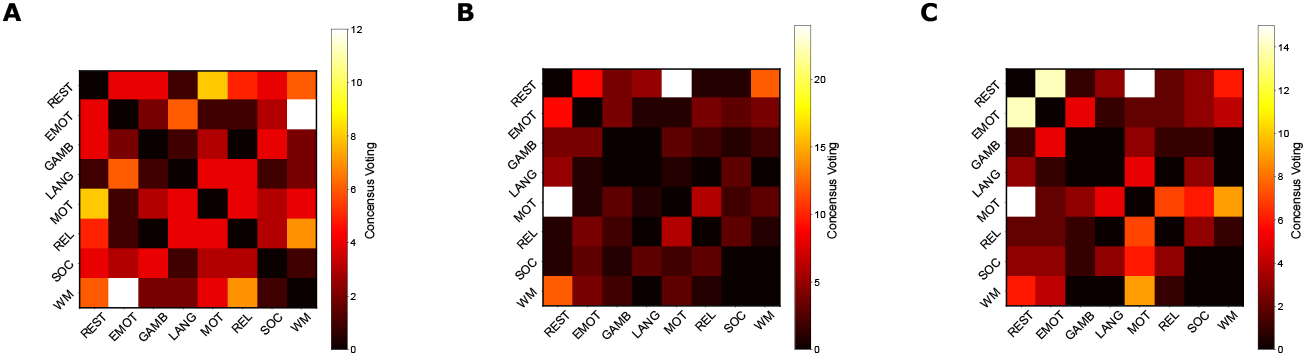
Individual consensus heatmap between tasks at the macroscopic level. Distance matrices between functional networks in 100 unrelated subjects were collected, and for each pair of functional networks, the frequency of it appearing as the most distinct pair among 100 subjects was counted, resulting in a majority voting heatmap for 3 homology groups (**Panel A** is the zeroth homology, **Panel B** is the second homology, and **Panel C** is the third homology). The number in the voting matrix represents the number of times the corresponding pair revealed the highest distance in one subject, and all numbers in one heatmap triangle should sum up to 100 for 100 subjects.

Particularly, the zeroth homology displayed the largest variability with the max count of the task pair being the smallest among the three homology groups, thus resulting in a more diffused pattern in the consensus voting heatmap (see **Figure** 6A). This serves as another explanation for the impact of the choice of graph representation on the zeroth homology analysis that it is relatively more varied. However, we also see the resting-motor task pair as one of the task pairs that have a high frequency at the individual level H0 results. Furthermore, the first homology still demonstrates the consistency with the group-averaged macroscopic level as well as consolidated level analysis, where it not only has the motor task-resting state as the most frequent task pair, but the max count is also the highest, indicating the robustness of the first homology in identifying brain connectivity pattern with different activities (see **Figure** 6B). The second homology also shows the motor task-resting state pair as the most frequent task pair, which further validates our findings shown above (see **Figure** 6C). The individual level analysis on the macroscopic level adds another layer to the group-averaged level analysis, where either the variability in the zeroth homology or the consistency in the first and second homology both further agree with the interpretation from previous sections.

#### 3.5.2 All-to-REST, mesoscopic analysis

At the individual mesoscopic level, the amount of analysis increased dramatically, with 100 individuals, 8 tasks, 8 functional networks, and 3 homological classes. In this case, it is difficult to analyze the distance between homology groups as we did at the group-averaged level. As validated in previous studies as well as our macroscopic level analysis, the resting state analysis tends to be the most distinct task compared to other tasks that include some activity engagement [5]. Therefore, we collected individual level all-to-REST distance and compared them across the functional network dimension and task dimension.

For the mesoscopic level in an all-to-REST setting, we picked three functional network pairs that have the highest distance measure from the group-averaged results (section 3.4.1) for all three homology groups. For each pair of functional networks, we collected 14 vectors, with each FN having 7 vectors containing 100 individual level distance measures between the 7 non-resting-state tasks and resting state task (see **Figure** S10), and then we compared the KL divergence between the two functional networks with vectors from the same non-resting-state task (**Figure** 7). In other words, the KL divergence measures the difference between two distributions (two functional networks respectively for all subjects) of the distance measure between the non-resting-state task and resting-state task.

**Figure 7.**
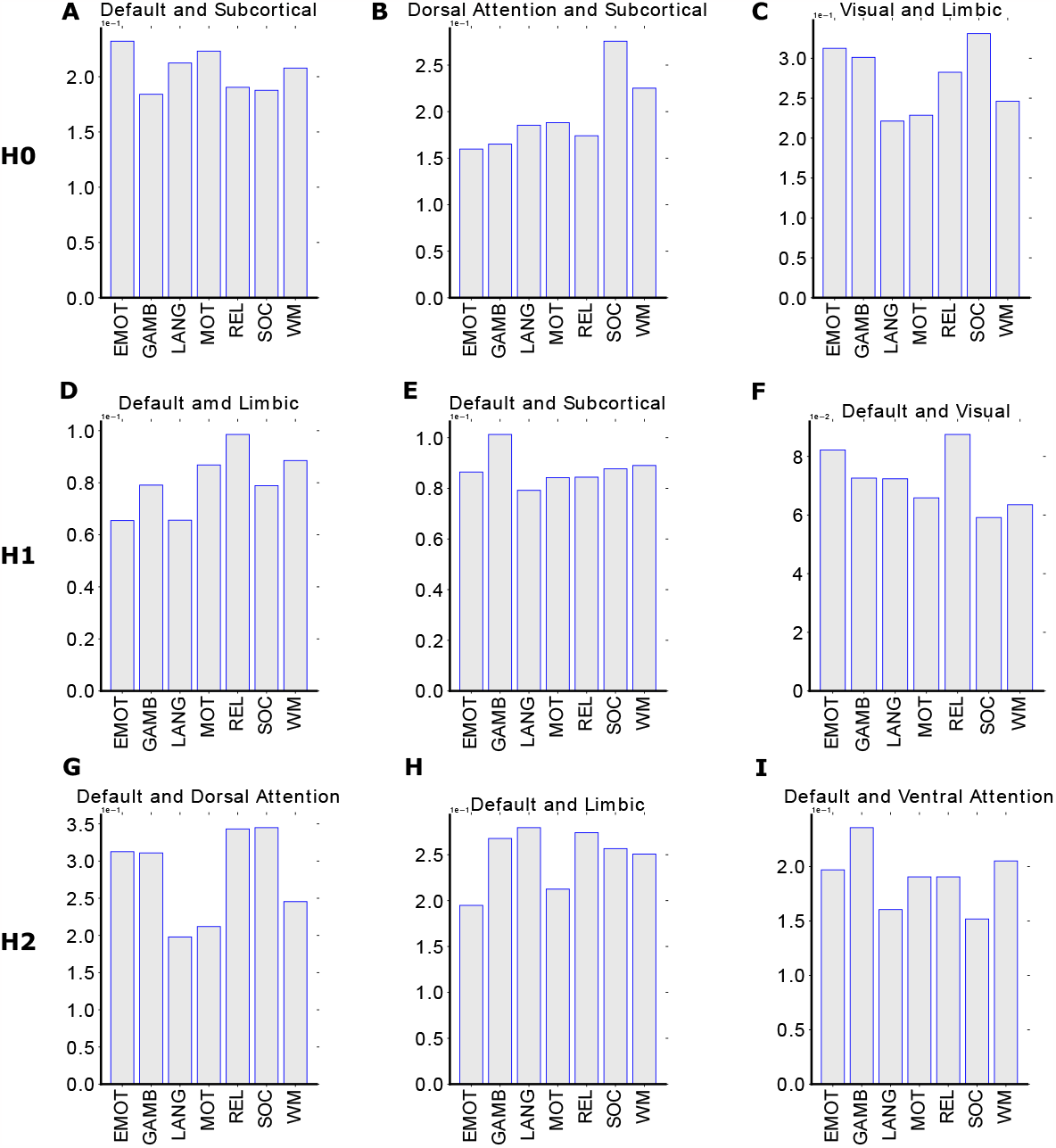
KL divergence plot for top three functional networks pair in all-to-RESTING setting. Rows represent homological groups (**Panel A-C** is the zeroth homology, **Panel D-F** is the first homology, and **Panel G-I** is the second homology) and each has three panels consisting of the top three most distinct pairs of functional networks inferred from the group-averaged mesoscopic analysis. The bar plot demonstrates the KL divergence between the selected pair of functional networks, in terms of the 100 individual-level distance between the resting state with other tasks.

For zeroth homology, we find that the social task is more differentiated from the resting state compared to others when we consider functional network pairs of dorsal attention and subcortical, as well as visual network with the limbic system (see **Figure** 7, panel B, C). These results take the consideration of both task activities and interactions between functional networks at the same time, indicating that the selected pair of functional networks have very different brain connectivity configurations in social tasks compared to the resting state. The default mode is still involved in the most selected pair of functional networks in the resting state, and the relational task has a very high KL divergence compared with the resting state in many functional pairs for the first and second homology, including default mode with limbic, subcortical, visual, dorsal and ventral attention (see **Figure** 7, panel D-I).

#### 3.5.3 All-to-REST, task analysis

The task analysis in an all-to-REST setting provided another perspective where the observation of functional network reconfiguration from resting state to other tasks is highlighted. In this case, we fixed the task that compared with the resting state and focused on the KL divergence between all pairs of functional networks in the first phonological order (see **Figure** 8, see **Figure** S11 and **Figure** S12 for the zeroth and second homology). To demonstrate the reconfiguration from resting to other tasks, we selected the top five largest KL divergences for each task and ranked them by the line strength in the circular plot.

**Figure 8.**
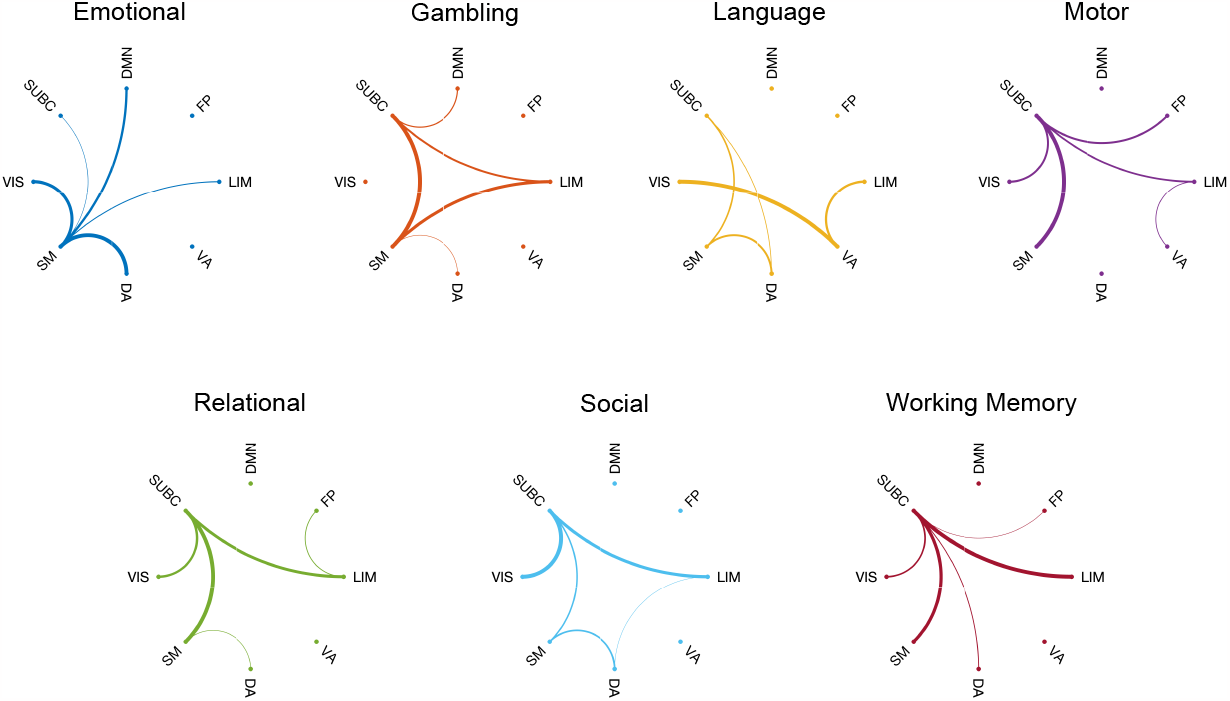
KL divergence circular plot for 7 fMRI tasks-to-RESTING with functional network comparison in H1. Here we fixed the task which compared with the resting state and visualized the top five KL divergence between functional networks. The KL divergence is normalized with regard to the top five measures and demonstrated by the strength of circular connectivity.

Some of the tasks displayed very unified patterns, such as the emotion task and working memory task, where all the highest KL divergence included one functional network (see **Figure** 8, emotion and working memory panel). The observation drawn from those two tasks showed that the reconfiguration from resting state to emotion task actually involves a lot of activities for somatomotor, and shifts to working memory task will require the subcortical region to take the most response. The somatomotor network includes most of the somatosensory area, which is closely related to emotional regulation, and the subcortical region is known to be involved in complex activities such as memory-related activities. In addition, we also observed that the somatomotor network also has the strongest link in the motor task, and the subcortical region is present as the dense connectivity hub in many task plots, which is an indication of the common underlying mechanism of brain circuit shifts from resting states to any other activities (see **Figure** 8, gambling, motor, relational, and social panel).

## 4 Discussion

At the heart of many complex systems resides a set of fine-tuned mesoscopic structures whose roles have been linked with complex orchestrations of emergent phenomena. Understanding complex higher-order behaviors arisen at a scale between the mesoscopic (brain regions) and macroscopic (whole-brain) level would set the stage to a more comprehensive understanding of the human brain large-scale functional circuitry. There are two kinds of mesoscopic structures: i) local/quasilocal (e.g., ground-truth communities) and ii) non-local such as topological strata of complex networks. In this work, we proposed a TDA formalism to disentangle the higher-order properties of brain sub-circuits (FNs) among different fMRI tasks. The major contributions of our framework on higher-order brain systems over other existing ones [4, 56, 61, 65] are that *i)* this framework allows the study of non-localized properties of an *a priori* set of localized/quasilocalized sub-networks, *ii)* through this innovative mesoscopic kernel proposal, we observed various results that align well with the current knowledge in network neuroscience and also highlighted the resting-state dichotomy of default mode network as well as the role of the limbic system in the process of functional (re)configuration, *iii)* we included not only within-task and within-FN scenarios, but also investigated the bi-level analysis that considered both task and FN levels at the same time. The construction of fMRI brain connectivity and Yeo’s ROI-to-FN mappings enabled multi-level homological group calculation and corresponding graph-based analysis. With 7 different tasks in addition to resting state, previous studies found that the brain functional reconfiguration in macroscopic (global-level) is hard to observe, while different tasks will rather trigger more shifts in mesoscopic structure (brain functional networks level) [20,24,48]. Hence, we organized our framework in 5 settings: a) group-averaged global level, group-averaged consolidated level, c) group-averaged mesoscopic level, d) individual global level, and e) individual all-to-REST level with functional network analysis and task analysis. At the first three levels, we conducted the topological data analysis at the group-representative level, which gives a broader view of the homological landscape between tasks and functional networks. When we look at the individual level (each subject’s FCs), we took a different approach from other existing brain connectivity fingerprint frameworks [1, 24]. Specifically, in the first step, we used consensus analysis to infer group-level behavior, as opposed to using simple averages. In the first step, we computed the distance measures on an individual basis by using the KL divergence to compare the distribution of individual-level distance. Through this setting, we found that three homological groups provided complementary insights in both task and subject domains. More specifically, the zeroth homology measures the connected components; the first homology measures the 2-dimensional hole encapsulated by one-dimensional functional edges; the second homology measures the 3-dimensional cavities encapsulated by 2-dimensional triangles. These homological groups and their algebraic structures are hypothesized in our paper to characterize topological spaces parameterized by the brain connectivity network.

Noticeably in work led by Fox and colleagues [32], the authors suggested that emotion task might be regulated by reduced functional activity attenuated by self-referential aspects of such task. In general, “harder” tasks (i.e., relational) require an increasing level of global integration which should reflect through a relatively small number of connected components (smaller Betti number 0). It is worthy to note that the motor cortex was identified as the hub of broadcasting transduction [5] which contains brain regions that are critical to broadcasting information to other regions of the brain. Compared to the resting state - the absence of cognitive requirement from fMRI tasks, motor task, which employs motor cortex brain regions, modulates global integrative cooperation among brain regions by forming first-order cycles across FNs. Combining both zeroth (connected components) and first homological (graph-theoretical cycles) distance results, we see that there exists a cognitive “switch” taking place at a global level to form connectivities that result in *i)* less number of connected components and *ii)* more globally integrated FNs as reflected by first-order cycles.

By consolidating the global view of the group-averaged connectome, we found that the H1 homology displayed stable topological invariants with its consistency in the most distinct pair of tasks as well as pertaining to a clear block diagonal structure on the distance heatmap. Both global and consolidated views displayed significant signals that the resting state and motor task are the most different task pairs [21, 42], while they are also the first and second distinct tasks in terms of the average distance (see **Figure** 4B,C). In this case, a simple observation we can draw from the analysis is that the brain takes some reconfiguration from resting state to other non-motor tasks, and then it requires further shifts in connectivity to get to the motor task. In addition, we further studied the individual-level homological scaffolds and performed group-level consensus voting on the most differentiated pair of tasks over 100 unrelated subjects (see **Figure** 6B,C). The H1 and H2 majority voting results again showed that the motor task is the most apart task from the resting state, and H1 also has the highest frequency count on the largest count among all three homological groups, indicating that it has the most consistent and robust capability to understand the homological scaffold in brain connectivity topological space.

Noticeably, the strong topological invariant of the H1 homology between the macroscopic (whole-brain) level with consolidated (super-graph) level demonstrated the existence of self-similarity property unraveled by the higher-order properties of brain functional sub-circuit [67, 69, 70]. Regarding the macroscopic level of the brain connectome as the “zoomed-in” representation of the consolidated graph, the overall pattern of the Wasserstein distance between tasks still holds. While both the macroscopic level and consolidated level have the resting-state task and motor task pair as the most differentiated task pair, further observation was found by looking at the row in the distance heatmap that involves resting state task and motor task all have high Wasserstein distance, together forming a block pattern that separates resting state task as well as motor task from the other tasks. This phenomenon guarantees the “scale-free” property of the first homological group on the complex brain system and provides a consistent potential for this topological framework for other higher-order complex network systems [67, 69].

We partitioned the brain connectome with the 7 Yeo functional network as well as a subcortical structure, resulting in 8 separate sub-networks. Since the resting state brain connectivity structure is the closest to Yeo’s partition, the first assessment that we did at the mesoscopic level was to fix the resting state task and compare the distance between two functional networks. The mesoscopic level analysis captured the “functional dichotomy” of the default mode network in the resting state by both the most differentiated task pairs as well as the highest average distance (see **Figure** 5B,C), where default mode is the most dominant network [24, 32, 40]. The brain network studies typically focus on either the within-task configuration or within-network configuration [5, 10, 12, 14, 61], the individual-level functional network partition further revealed patterns in the brain that are shifted between resting state to other tasks as well as between two functional networks. The individual all-to-rest mesoscopic analysis considered both task and functional network “switches”. Such bi-level perspective allows the investigation of the most distinct functional network pairs in resting state on their reconfiguration from resting state to other tasks (see **Figure** 7). While maintaining the bi-level design of the experiments, we flipped the two-level in the all-to-rest task analysis to investigate, from resting state to each task, how pairs of functional network are shifted (see **Figure** 8). The unique patterns in the top 5 pairs of functional networks also enabled hub identification in the process of the task switch, and closer tasks also displayed similar patterns, indicating that they underwent similar reconfiguration from the resting state.

This study has certain limitations. In the consolidation process from the global-level graph, we specifically opted for max normalization to construct the super graph. Since altering the normalization method may potentially modify the inter-connectivity of functional networks, future research could investigate different normalization techniques. For instance, using average connectivity to define the consolidated graph might impact not only the topological structure of the super graph but also its self-similarity properties from the homological kernels. Moreover, not only does the choice of the homological group influence the distance measure between tasks or functional networks, but the graph itself also plays a crucial role. Our experiments were solely conducted on the Glasser parcellation with 374 nodes (360 cortical regions + 14 sub-cortical regions). Exploring alternative parcellations in both brain cortical and subcortical regions ([58, 72]) and incorporating multiple parcellation scales could offer additional insights into mesoscopic cognitive reconfiguration and its scaling-related properties.

In summary, we presented a novel framework that uses persistent homology to characterize brain connectivity in the topological space. Based on the nature of each homological group, we selected different distance measures correspondingly. The zeroth persistent homology is all born at 0 so the Wasserstein distance is not a good fit, but the Haursdorff distance is more appropriate for measuring the 1D distribution of the point cloud. However, the first and second homology are closer to the diagonal in the persistent homology diagram, and thus the Wasserstein distance with partial mapping which serves as a simulation of moving one distribution to another in a geodesic setting would become better in this case. We validated that the first homology gives very consistent and topological invariant findings in different levels of analysis, which offers a scaling invariant perspective, and we find that the framework is capable of capturing signals that are well-studied in the literature, but also discovered some unique patterns in the brain circuit triggering diverse processes among different fMRI tasks and resting conditions. From a wider perspective, our formalism can be applied, beyond brain connectomics, to study non-localized coordination patterns induced by localized, pre-defined structures stretching across different complex network fibers.

## Acknowledgements

Data were provided (in part) by the Human Connectome Project, WU-Minn Consortium (principal investigators: David Van Essen and Kamil Ugurbil; 1U54MH091657) funded by the 16 NIH Institutes and Centers that support the NIH Blueprint for Neuroscience Research; and by the McDonnell Center for Systems Neuroscience at Washington University.

## Author Contributions

Duy Duong-Tran: Conceptualization; Formal analysis; Investigation; Methodology; Writing original draft. Ralph Kaufmann: Methodology; Investigation; Writing original draft. Jiong Chen: Investigation (Computation); Formal Analysis; Visualization; Writing original draft. Xuan Wang, Sumita Garai, Frederick Xu, Alan D. Kaplan, Yize Zhao: Writing original draft. Jingxuan Bao: Visualization. Giovanni Petri: Methodology; Writing original draft. Enrico Amico, Joaquin Goñi: Data curation; Writing original draft. Li Shen: Conceptualization; Formal analysis; Writing original draft; Project Supervision; Funding acquisition.

## Supplementary Material

### Persistent diagram

**Figure S1.**
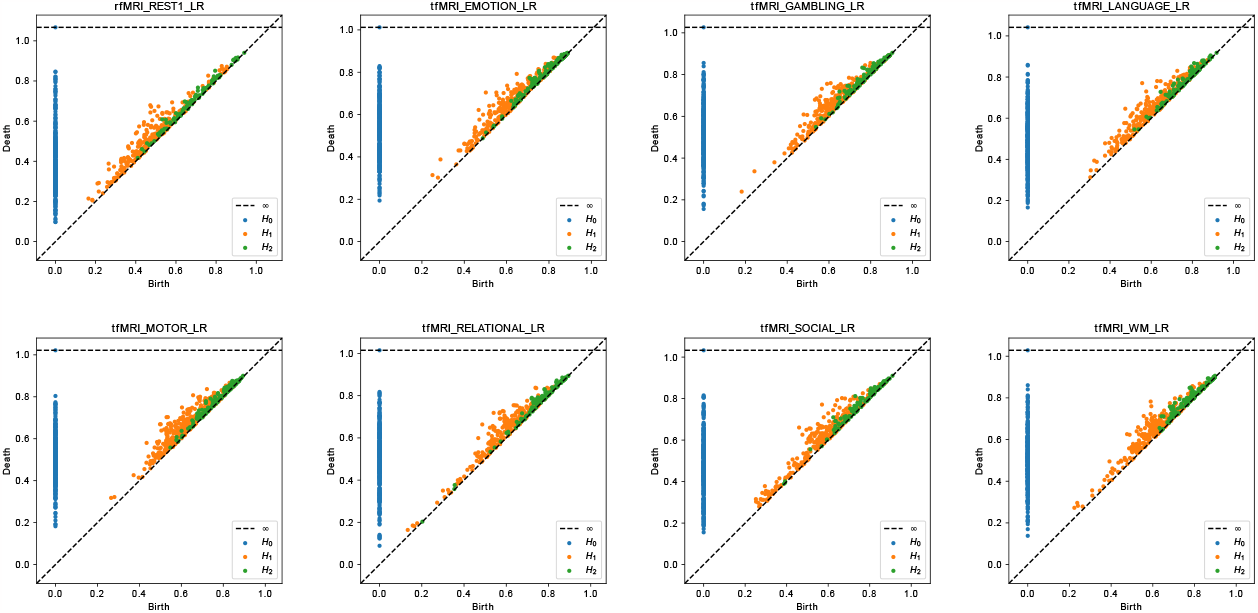
Persistent diagram of the global, group-averaged connectome. Figures shown above are the persistent diagram for all 8 tasks in 3 homological groups, represented by blue, orange, and green dots respectively. The connectome is obtained by taking the average of 100 unrelated HCP subjects’ functional connectome.

**Figure S2.**
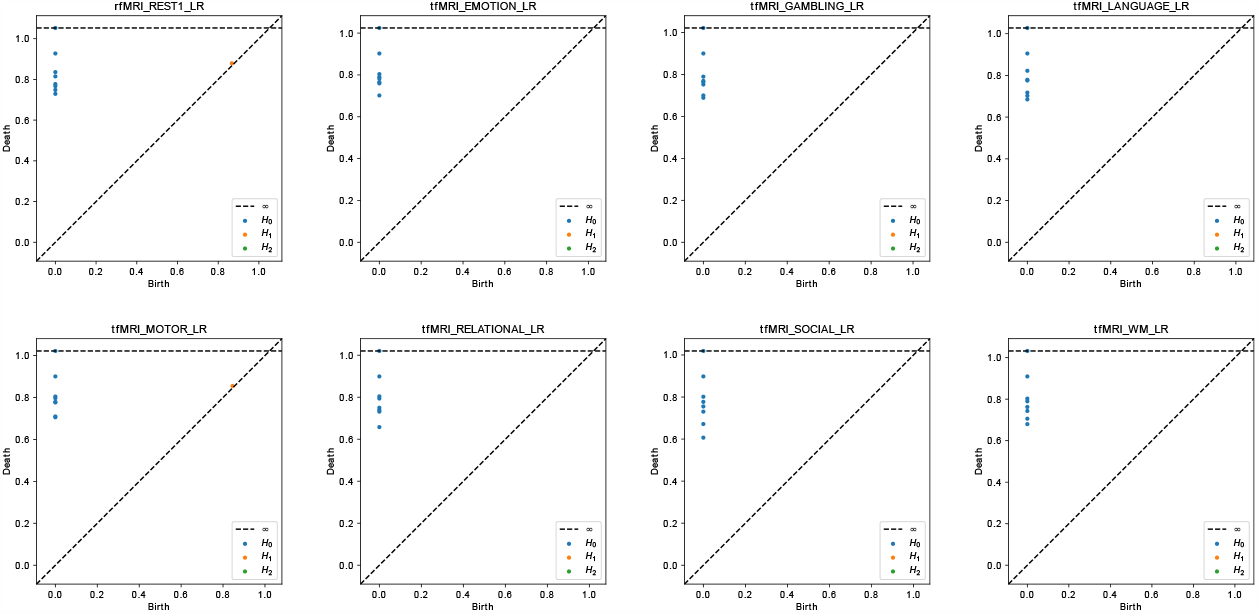
Persistent diagram of the consolidated, group-averaged connectome. Figures shown above are the persistent diagram for all 8 tasks in 3 homological groups. The connectome is obtained by consolidating the communities in the global connectome by Yeo functional networks. Since the consolidated graph only has 8 nodes, there are only two homological groups exist (the zeroth homology represented by blue dots and the first homology represented by orange dots).

### Group-mesoscopic level analysis for other tasks

**Figure S3.**
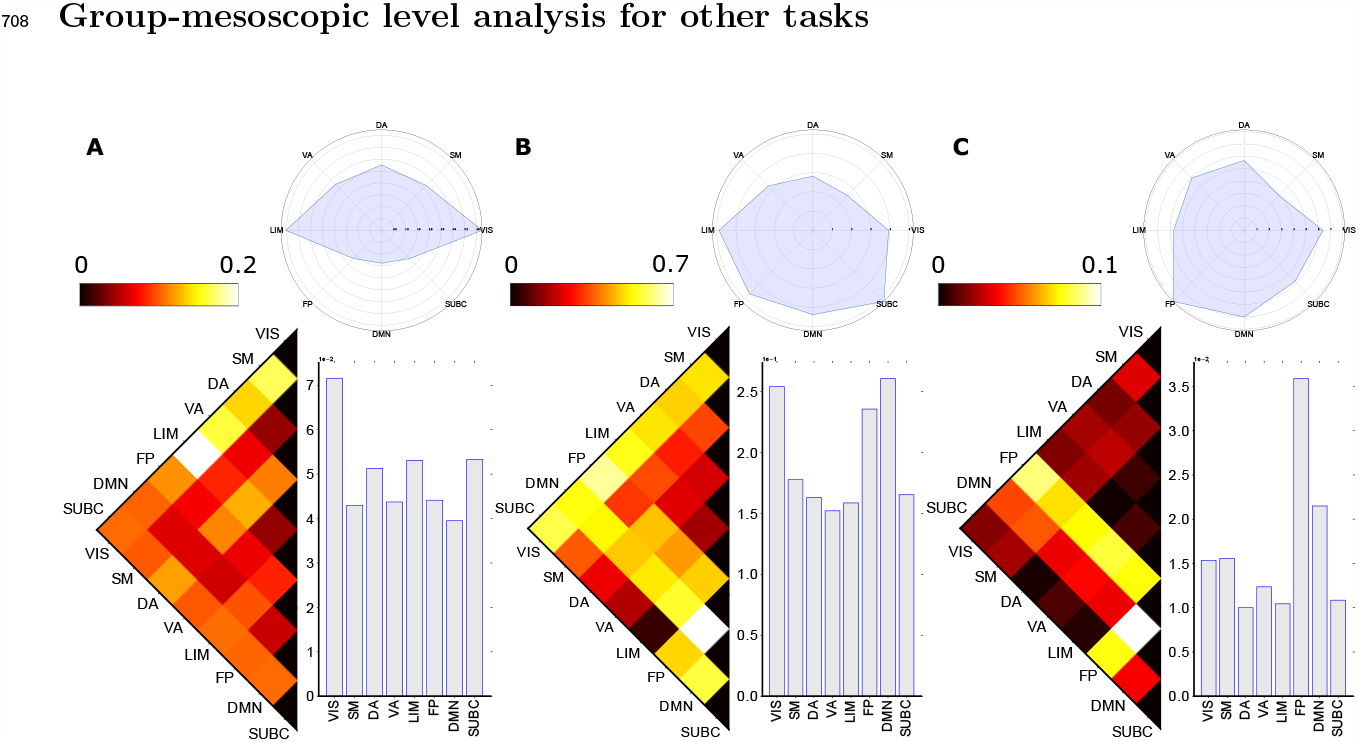
Group-Average Homological distances between brain circuits (FNs) in emotion task. In emotion task, the limbic-visual network pair is the most differentiated FN pair in zeroth homology, while the frontoparietal network and default mode network are the most distinct pair of FNs. The visual network and frontoparietal network have the highest average distance compared with other FNs in the zeroth and second homology, but in the first homology, the visual network, frontoparietal network, and default mode network all have high average distance. The limbic and visual networks also have high variance in distance compared to others.

**Figure S4.**
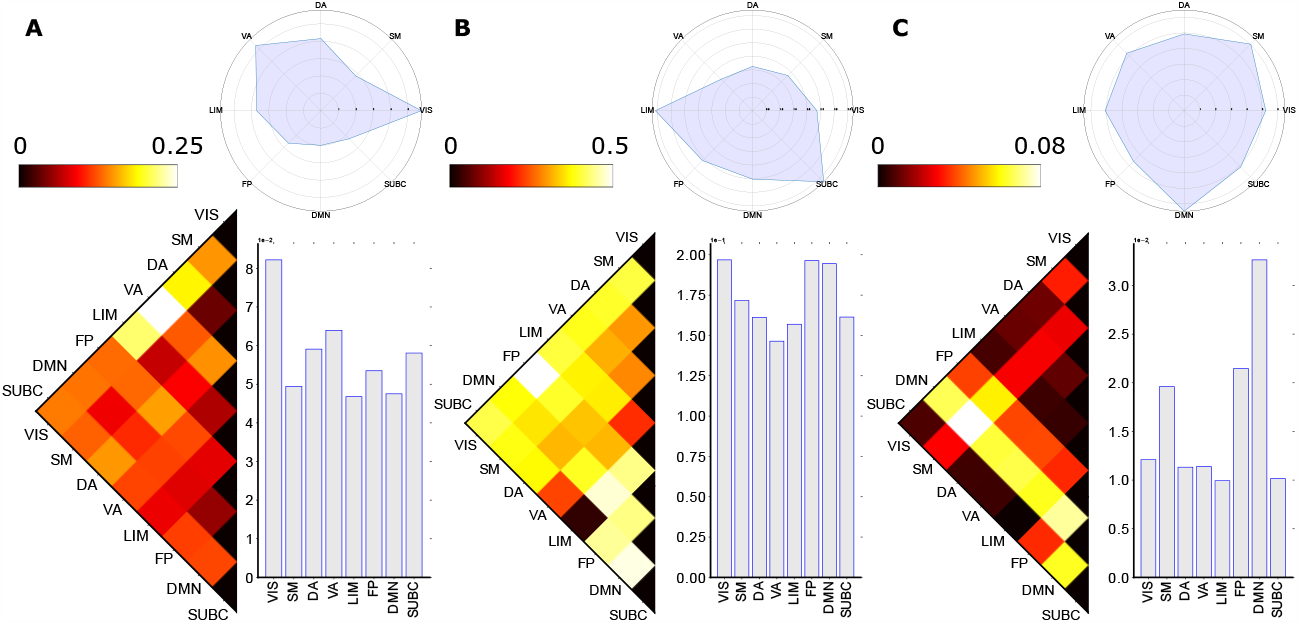
Group-Average Homological distances between brain circuits (FNs) in gambling task. In gambling task, the visual network with ventral attention pair, visual network with frontoparietal network pair, and somatomotor network with default mode network pair are the most differentiated FN pairs in the zeroth, first, and second homological group respectively. The visual and default mode networks have the highest average distance in the zeroth and second homology, while in the first homology, the visual network, frontoparietal, and default mode network all have high average distance. The ventral attention and visual networks have high variance in zeroth homology, and the limbic and subcortical systems have high variance in the first homology.

**Figure S5.**
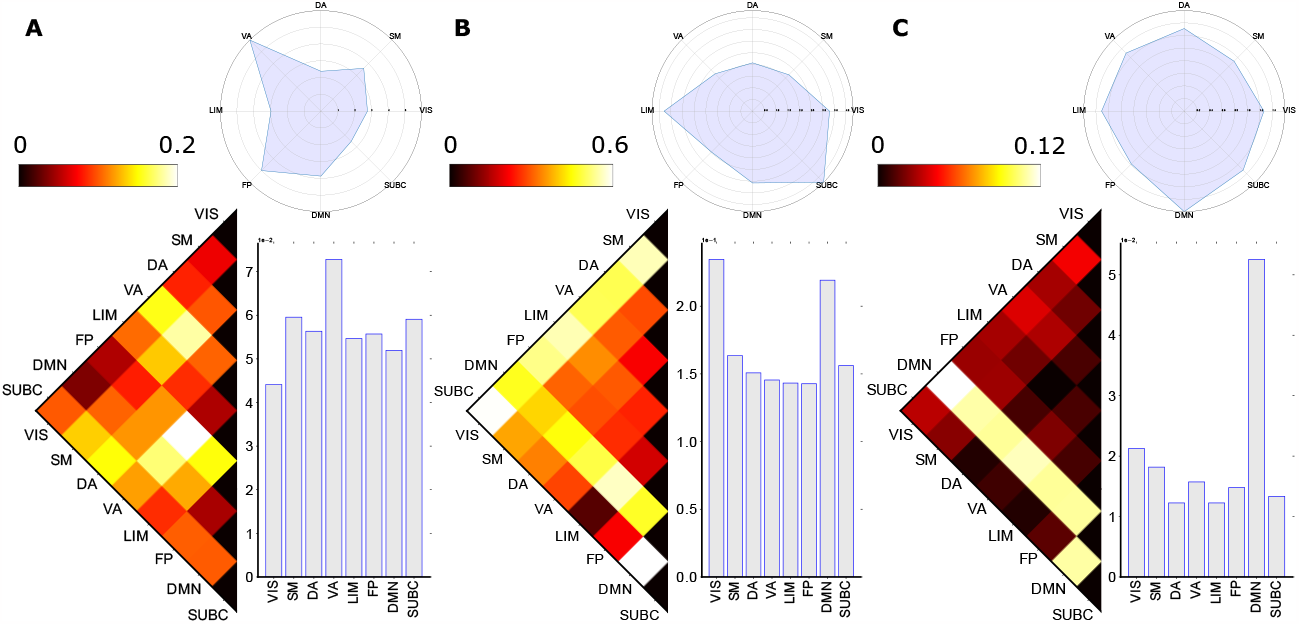
Group-Average Homological distances between brain circuits (FNs) in language task. In language task, the frontoparietal with ventral attention networks are the most distinct pair of tasks in the zeroth homology. In the first homology, the subcortical system with both visual and default mode networks all have a high Wasserstein distance. In addition, the default mode with visual networks is the most distinct pair of tasks in the second homology. The ventral attention, visual, and default mode networks have the highest average distance in the zeroth, first, and second homological groups, with ventral attention, subcortical, and default mode systems having the highest variance. The default mode also serves as the clear winner in the second homology in distance heatmap as well as average distance, indicating its high involvement in the language task.

**Figure S6.**
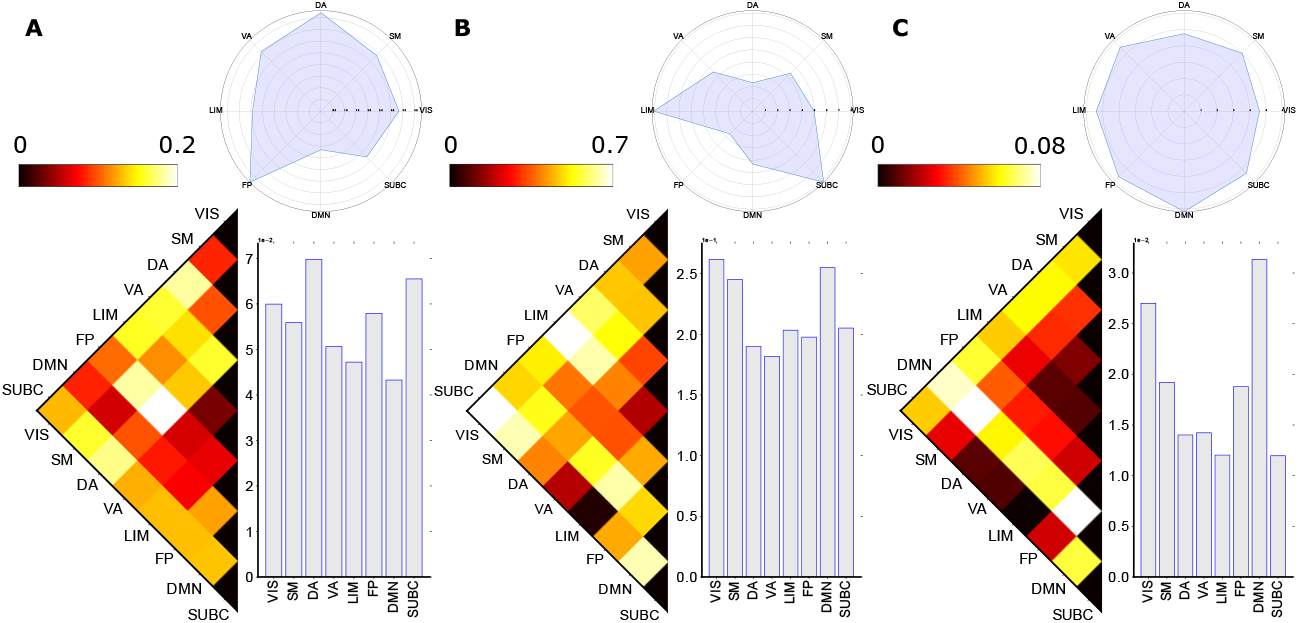
Group-Average Homological distances between brain circuits (FNs) in motion task. In motion task, the frontoparietal with dorsal attention networks are the most distinct pair of tasks in the zeroth homoloyg. In the first homology, the visual network with both subcortical and limbic system all have a high Wasserstein distance. In the second homology, the default mode with both somatomotor and frontoparietal networks have a high Wasserstein distance. The dorsal attention, visual, and default mode networks have the highest average distance in the zeroth, first and second homological groups. The visual network and frontoparietal networks have the highest variance in the zeroth and second homology, while in the first homology, the default mode network and subcortical system both have a high variance.

**Figure S7.**
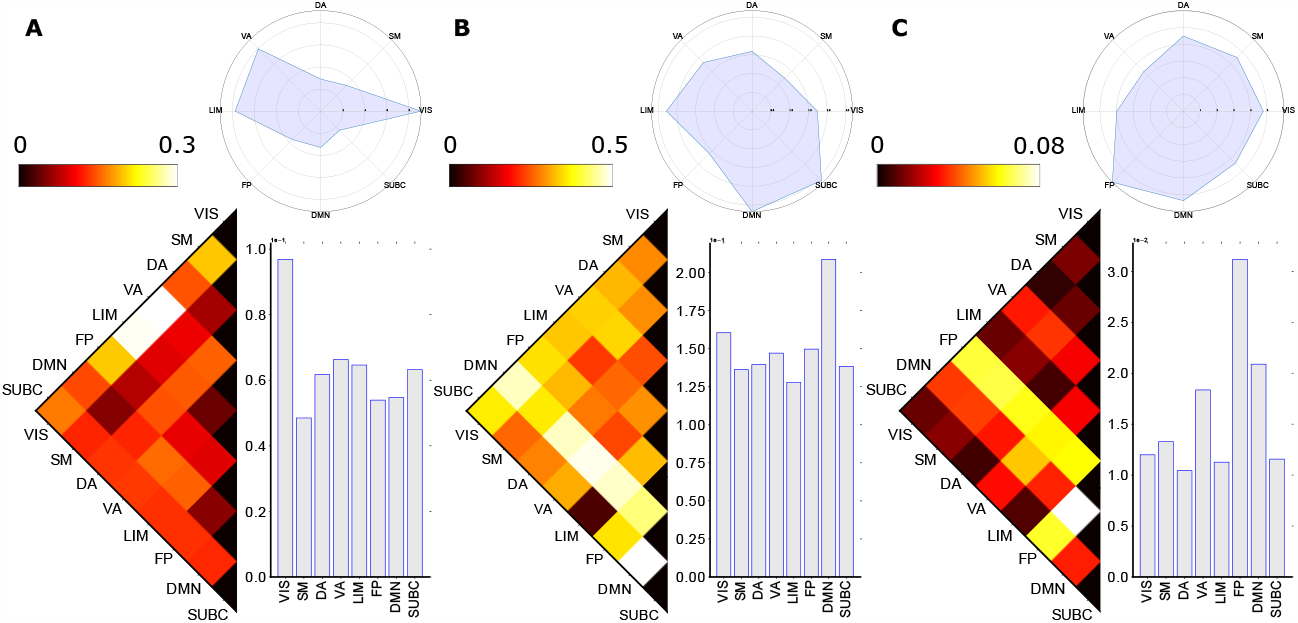
Group-Average Homological distances between brain circuits (FNs) in relational task. In relational task, the visual network with both limbic and ventral attention networks have the highest Hausdorff distance. The visual network with ventral attention also has the highest Wasserstein distance in the first homology, and in the second homology, the frontoparietal and default mode network pair has the highest distance. The relational task is also consistent in the average distance with the zeroth and first homology, having the visual network as the most distinct network. The frontoparietal network has the highest average distance in the second homology. In all homological groups, the visual network has very high variance, and the subcortical system in H1 and the frontoparietal network in H2 also have high variance compared to others.

**Figure S8.**
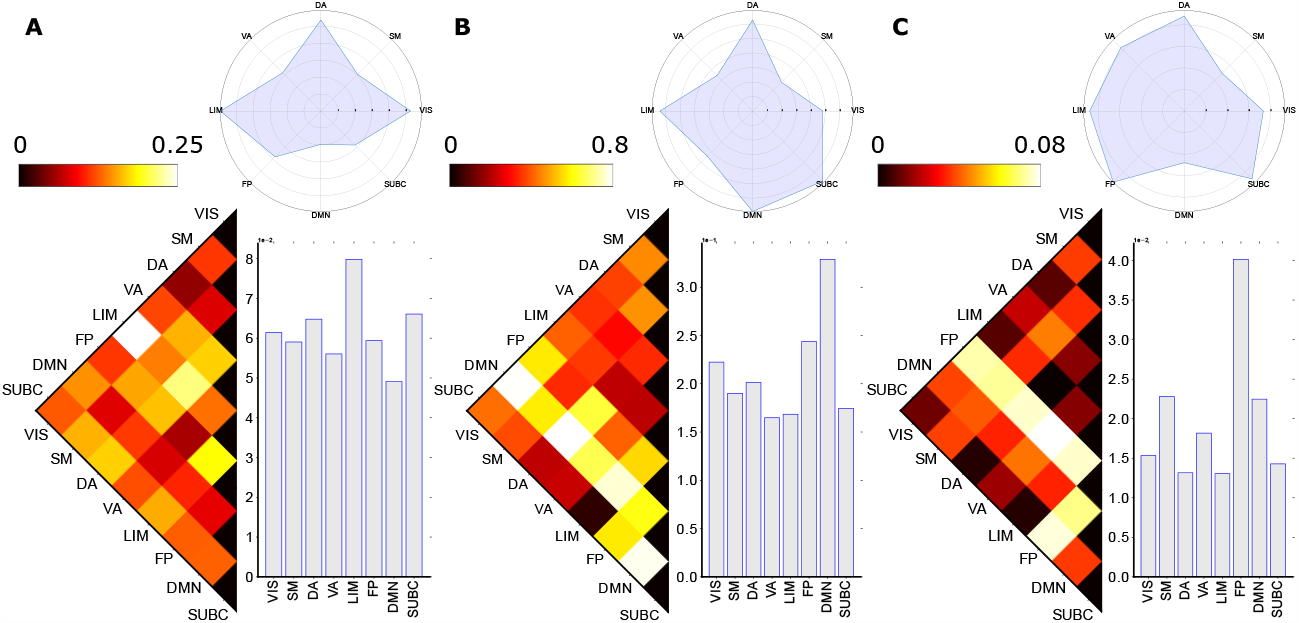
Group-Average Homological distances between brain circuits (FNs) in social task. In social task, the limbic network and visual network pair is the most differentiated FN pair in the zeroth homology. The first homology has multiple distinct FN pairs, and all of them involve the default mode network. A similar pattern is observed in the second homology with the frontoparietal network as the consistent network in those FN pairs. These results also correspond with the average distance, where the limbic network, the default mode network, and the frontoparietal network is the most distinct network in the three homological groups, and at the same time they also have the highest variance.

**Figure S9.**
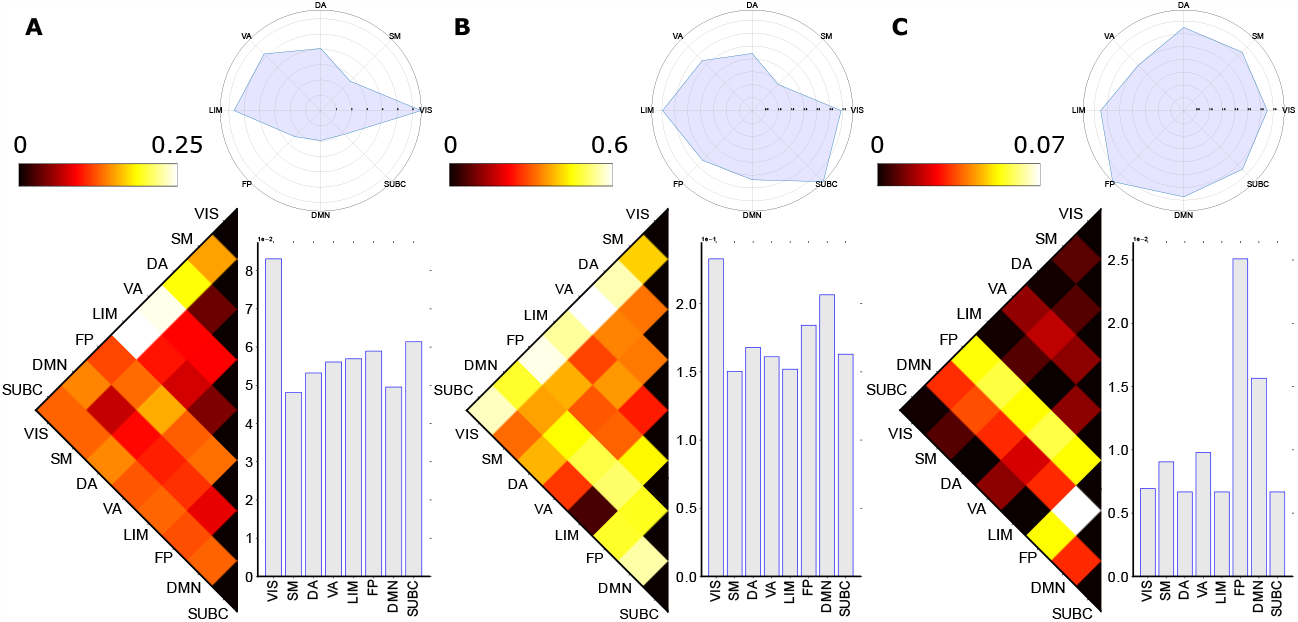
Group-Average Homological distances between brain circuits (FNs) at working memory task. In working memory task, the zeroth homological group has the limbic-visual network pair as the most differentiated pair, and the visual network also has the highest average distance and variance. The ventral attention and visual network pair has the highest distance in the first homology, with the visual network still being the most distinct network, but the subcortical system has the highest variance. In the second homology, the strong separation in the frontoparietal network is consistent with the average distance plot as well as the variance plot that the frontoparietal network is the most differentiated FN, and particularly with the default mode network, the FN pair has the highest Wasserstein distance.

### Individual-mesoscopic level analysis

**Figure S10.**
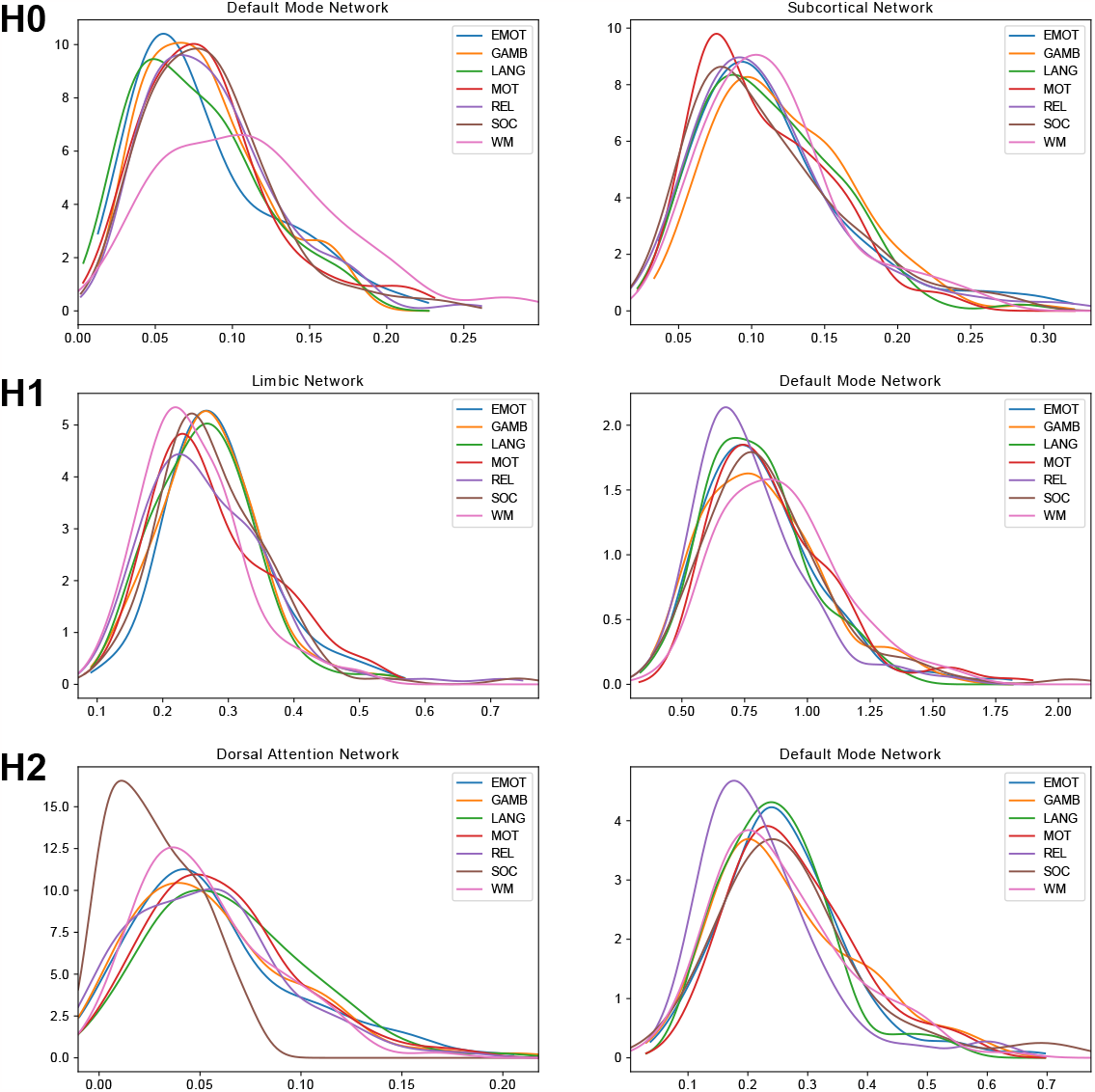
Distribution of pairs of functional networks having the highest distance in three homological groups. The top row (H0), middle row (H1), and bottom row (H2) show the distribution of distance measures between the resting state and the other 7 tasks. Each row involved the FN pair with the highest distance in the group-level mesoscopic analysis at the resting state. The distribution is used to calculate the KL divergence described in the main article.

**Figure S11.**
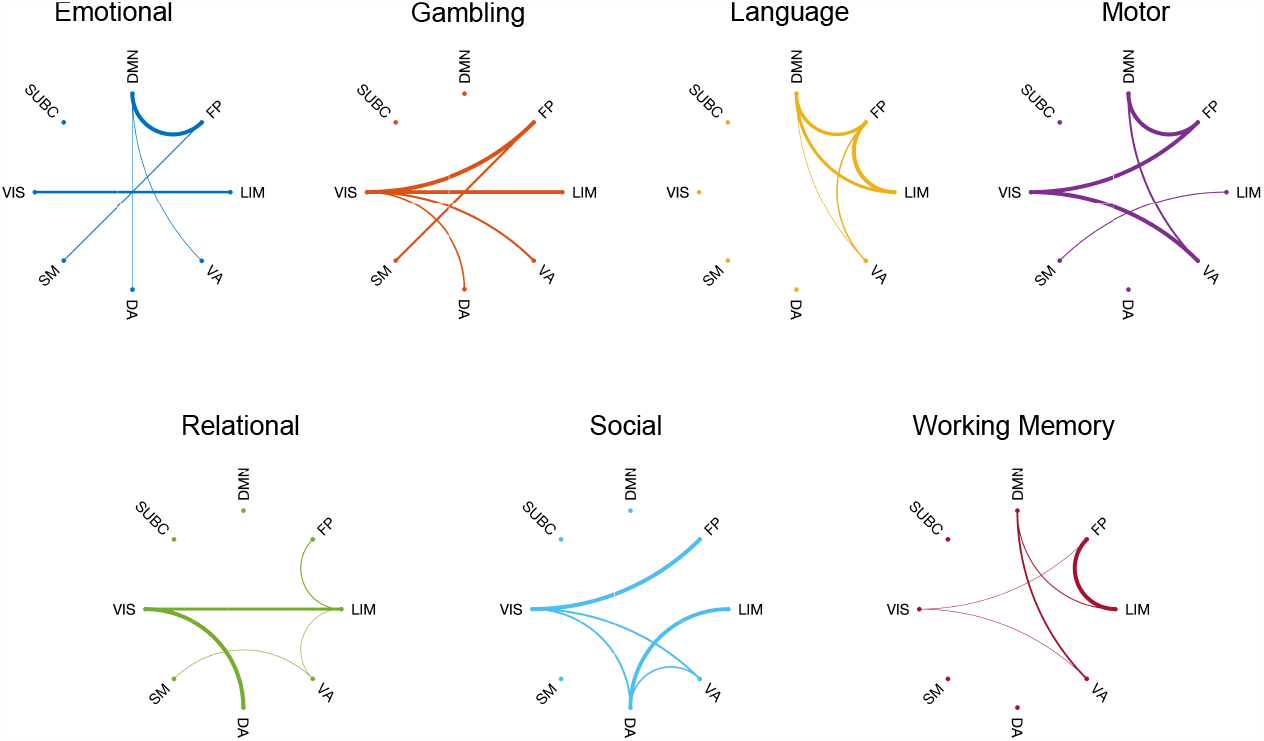
KL divergence circular plot for 7 fMRI tasks-to-REST with functional network comparison in H0. The KL divergence between functional networks (see **Figure** S10 for example) is visualized with the task-wise circular plot. The top five KL divergence is normalized and demonstrated by the strength of circular connectivity. The pattern in the zeroth homology is more heterogeneous, but we can still identify some hubs such as the visual network in the gambling task.

**Figure S12.**
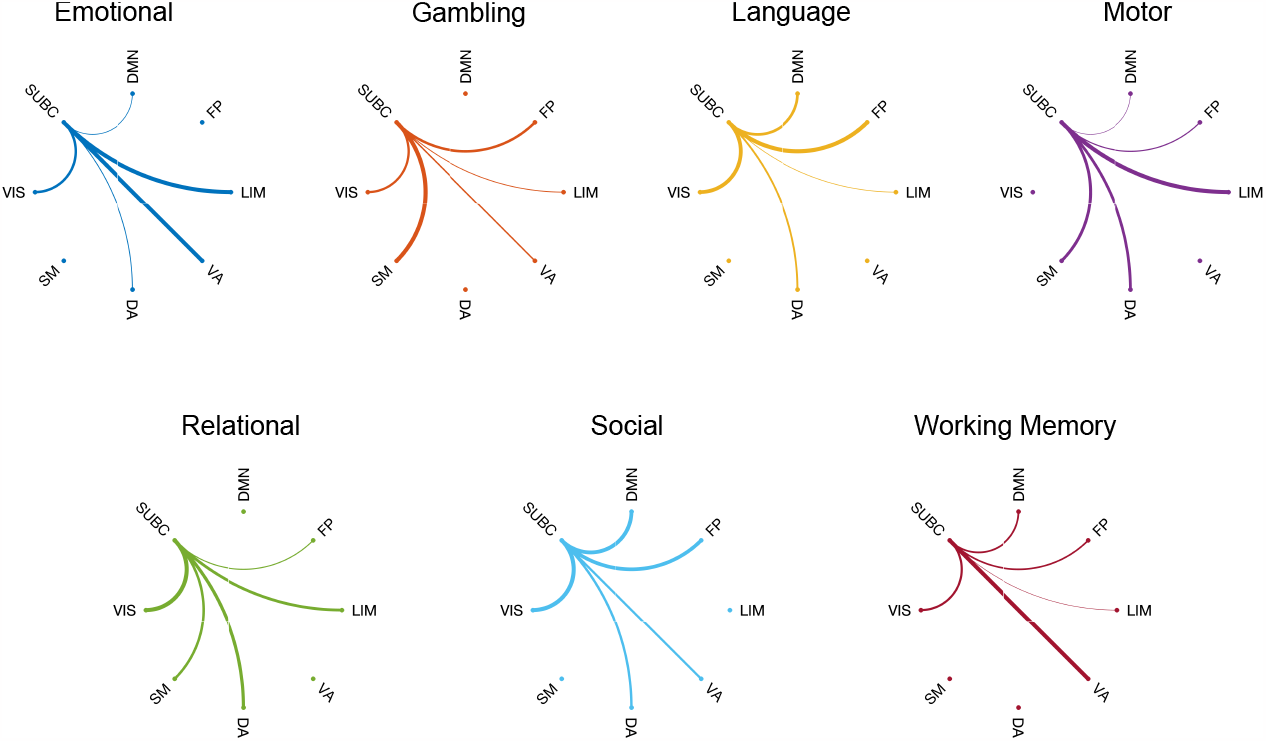
KL divergence circular plot for 7 fMRI tasks-to-REST with functional network comparison in H2. The highest five KL divergence is visualized with the task-wise circular plot and normalized for clearer demonstration. The H2 homology provides a higher level of topological structure view, where we can find that the subcortical system is mostly the hub for all tasks.

## Footnotes

1 https://www.humanconnectome.org/storage/app/media/documentation/s1200/HCP_S1200_Release_Reference_Manual.pdf.

2 http://www.humanconnectome.org/software/connectomeworkbench.html

